# Adaptation to ARF6-depletion in KRAS-driven PDAC is abolished by targeting TLR2

**DOI:** 10.1101/2023.11.30.569405

**Authors:** Ritobrata Ghose, Shubhamay Das, Julia Urgel-Solas, Fabio Pezzano, Debayan Datta, Upamanyu Ghose, Antoni Ganez Zapater, Natalia Pardo Lorente, Lorena Espinar Calvo, Laura Garcia, Chiara Cannata, Verena Ruprecht, Ana Janic, Sara Sdelci

## Abstract

Metastasis is responsible for nearly 90% of all cancer-related deaths. Despite global efforts to prevent aggressive tumours, cancers such as pancreatic ductal adenocarcinoma (PDAC) are poorly diagnosed in the primary stage, resulting in lethal metastatic disease. RAS mutations are known to promote tumour spread, with mutant KRAS present in up to 90% of cases. Until recently, mutant KRAS remained untargeted and, despite the recent development of inhibitors, results show that tumour cells develop resistance. Another strategy for targeting mutant KRAS-dependent PDAC proliferation and metastasis may come from targeting the downstream effectors of KRAS. One such axis, which controls tumour proliferation, invasiveness and immune evasion, is represented by ARF6-ASAP1. Here we show that targeting ARF6 results in adaptive rewiring that can restore proliferation and invasion potential over time. Using time-series RNA and ATAC sequencing approaches, we identified TLR-dependent NFκB, TNFα and hypoxia signalling as key drivers of adaptation in ARF6-depleted KRAS-dependent PDAC. Using in vitro and in vivo assays, we show that knocking down TLR2 with ARF6 significantly reduces proliferation, migration and invasion. Taken together, our data shed light on a novel co-targeting strategy with the therapeutic potential to counteract PDAC proliferation and metastasis.

**GRAPHICAL SUMMARY:** 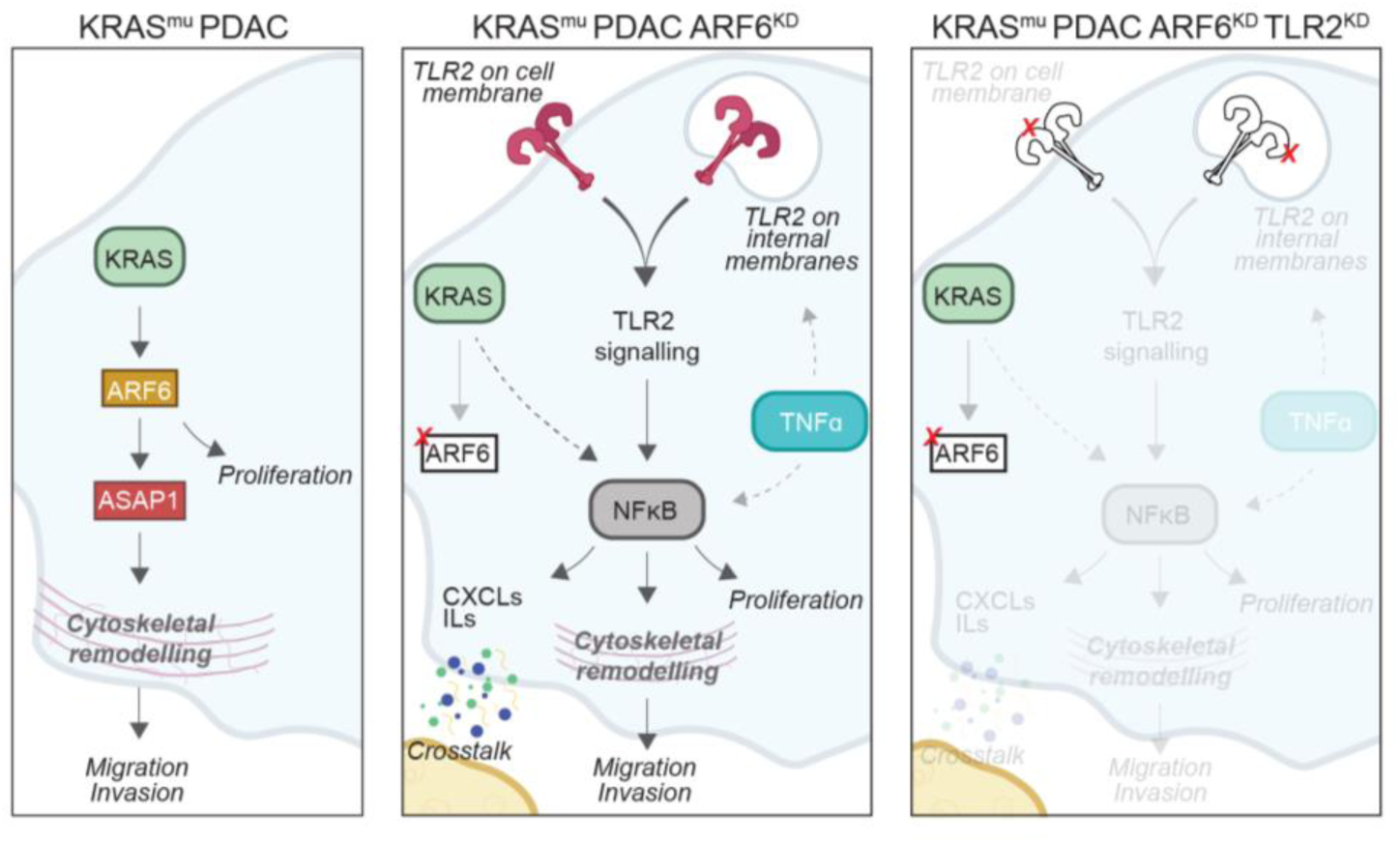

## INTRODUCTION

Pancreatic cancer is an intractable disease and the 7^th^ leading cause of cancer mortality worldwide, 3^rd^ in the USA and 4^th^ in Europe.^1^ Although incidence rates are lower compared to other cancer types, it is the disproportionately high mortality rate however, that makes this cancer so deadly. KRAS is the most commonly mutated of the RAS oncogenes and is most prevalent in pancreatic ductal adenocarcinoma (PDAC),^2^ accounting for between 69% to 95% of all PDACs.^3^ In the majority of cases, KRAS mutations are single-base missense mutations, 98% of which are at Glycine 12 (G12, accounts for 81%), Glycine 13 (G13, accounts for 14%) or Glutamine 61 (Q61, accounts for 2-3%).^4^ KRAS, which is one of the main drivers of PDAC has remained undruggable until very recently. AMG510, also known as sotorasib^5^ was recently found as a KRAS G12C inhibitor and is now the only FDA-approved drug in clinical trials against KRAS G12C. Other RAS inhibitors in the pipeline include KRAS G12C inhibitor MRTX849,^6^ KRAS G12D inhibitor MRTX1133,^7^ and HRAS G60A inhibitor NSC290956.^8^ Finally, a recent publication brought to light the pan-KRAS inhibitor BI-2865, which does not target either HRAS or NRAS.^9^ BI-2865 was able to bind to wild-type and mutant KRAS with high affinity, preventing its further activation. While finally having KRAS inhibitors is exciting, their success remains to be tested in clinics. A recent preliminary result from the AMG510 clinical trial warrants caution in its application.^10^ A comparison of the pre-treatment and post-treatment genomic landscape of patients treated with AMG510 identified a series of acquired mutations in a subset of patients, which had the ability to induce drug resistance. They identified additional mutations in KRAS, NRAS, B-RAF, muscle (M)RAS, EGFR, fibroblast growth factor receptor (FGFR) 2 and the MYC proto-oncogene. Moreover, these activating mutations of the various RAS proteins results in reactivation of the KRAS-downstream signalling axes given by the MAPK pathway resulting in drug resistance and reversion to tumour progression.^10^

Another strategy of targeting KRAS-dependent PDAC is the targeting of important downstream signalling factors, altogether limiting the key pro-tumorigenic effects. One study attempted the inhibition of the two mTOR complexes (mTORC1/2) in PDAC, to overcome KRAS and MEK inhibitor resistance.^11^ When inhibiting both KRAS and mTORC1/2, a synergistic reduction in cell survival and tumour proliferation was achieved. Nevertheless, ERK activation was unfavourably altered,^11^ bringing into question whether the targeting of KRAS itself may result in pro-tumorigenic alterations as a result of tumour adaptation. Another recent study investigated the synergistic downregulation of MEKs using Trametinib and epigenetic modulators of the BET family using JQ1.^12^ In PDAC, the combination of Trametinib and JQ1 synergistically reduced cell proliferation and induced cell death.

Overall, it becomes evident that targeting KRAS or master regulators by themselves is neither easy and can be challenging in terms of conclusive results. Another strategy which may be more viable is the targeting of downstream factors that are critical enough to regulate major pathways. In that context, understanding the time-dependent landscape of PDACs which are subjected to perturbations at different hierarchical levels of important signalling cascades will provide insight into the potential adaptation mechanisms which will yield robust synergistic anti-tumour effects. One such axis is given by ARF6-ASAP1 signalling. ARF6 is a small GTPase of the Ras family of GTPases and is a major mediator of KRAS signalling. Loss of ARF6 has been studied to impair proliferation, migration and invasion,^13–19^ which in turn has prompted the development of ARF6 inhibitor NAV-2729,^20^ and is considered as a potential therapeutic target.^21^ GTPases use effector proteins for their downstream signalling. Among others, ASAP1 (also AMAP1 or DDEF1), is a major downstream effector of ARF6 in various cancers, but particularly important in PDAC.^15^ KRAS-downstream signalling acts on both ARF6 and ASAP1 to downregulate E-cadherin and promote a more mesenchymal phenotype.^13,15^ Additionally ASAP1 can also regulate cytoskeletal features such as β1 integrins leading to a PDAC invasive phenotype.^22^ In addition to promoting tumour invasiveness, ARF6, through ASAP1 also promotes the intracellular recycling and cell surface expression of programmed death ligand 1 (PD-L1), which binds to the PD-1 receptors on immune cells, leading to immune checkpoint evasion.^15^

Here, we perturbed KRAS-downstream ARF6 or ASAP1 in PDAC. For the first time, we show that downregulating ARF6, but not ASAP1 leads to an adaptive response in PDAC cells leading to the re-acquisition of proliferation and invasion phenotypes. By further performing time-dependent transcriptomic and chromatin accessibility analysis, we show that this adaptation may be attributed to a TLR2-mediated response together with the activation of NFkB, TNFa and Hypoxia pathways. Finally, we show that the co-knockdown of ARF6 and TLR2 almost abolishes tumour growth, and heavily deters metastatic potential of PDAC in vivo.

## RESULTS

### PDAC adapts to ARF6, but not ASAP1 loss

To understand the adaptive landscape of PDAC when KRAS-downstream signalling nodes at different hierarchical levels are perturbed, we treated MIA PaCa-2 cells with shRNAs targeting either ARF6 or ASAP1, and monitored them for a prolonged period of up to 21 days. Given that, ARF6 is a major regulator of KRAS signalling, and in general, various PDAC-related pathways, our readout from ARF6 will act as a proxy for an intermediate level factor in KRAS-downstream signalling. Conversely, ASAP1, which is an effector molecule of ARF6 signalling, will act as a proxy for a KRAS-signalling end point.

MIA PaCa-2 cells were transduced with lentiviruses containing shRNAs for either ARF6 or ASAP1 (Figure 1 A). A non targetting (NT) vector was used as control. Dowregulation of ASAP1 and ARF6 was checked by Western Blot at each time point (Supplementary Figure S1 A-B). Both ARF6 and ASAP1 have been strongly linked to proliferation and invasion of PDAC.^15^ To test the effect of ARF6 and ASAP1 loss on KRAS-dependent PDAC, we set up a series of time-dependent experiments (Figure 1 A). First, we performed Matrigel invasion assays using Transwells and MTT-based proliferation assays, taking measurements at the four time points – day 3, 7, 14 and 21 post-downregulation (Figure 1 A). As previously observed,^15^ treatment with shARF6 or shASAP1 resulted in an immediate decrease in invasion potential of MIA PaCa-2 cells in comparison to cells treated with shNT (Figure 1 B). We observed a significant decrease by 25% and 35% respectively for KD of ARF6 and ASAP1 on day 3. shASAP1-treated cells continued to exhibit significantly lower invasion by about 54%, 65% and 52% respectively on days 7, 14 and 21. Contrastingly, shARF6-treated cells showed an adaptive behaviour. They recovered their invasion capacity at day 7, which stabilised over time (Figure 1 B). We also observed similar trends in shARF6-treated PDAC cell line Panc-1 and colorectal carcinoma cell line HCT-116 (Supplementary Figure S1 C). Like invasion, we observed significantly reduced proliferation in shARF6 and shASAP1 treated cells (Figure 1 C). On day 3, shARF6 showed a proliferation rate that was about 48% less than shNT, and shASAP1 showed a proliferation rate which was about 66% less than shNT. shASAP1-treated cells continued to show much lower proliferation rates compared to shNT throughout the following three time points, being at approximately 39%, 59% and 54% that of shNT respectively. In contrast, and as in the case of invasion, shARF6-treated cells recovered over time, reaching up to shNT proliferation rates by day 21 (Figure 1 C).

**Figure 1.**
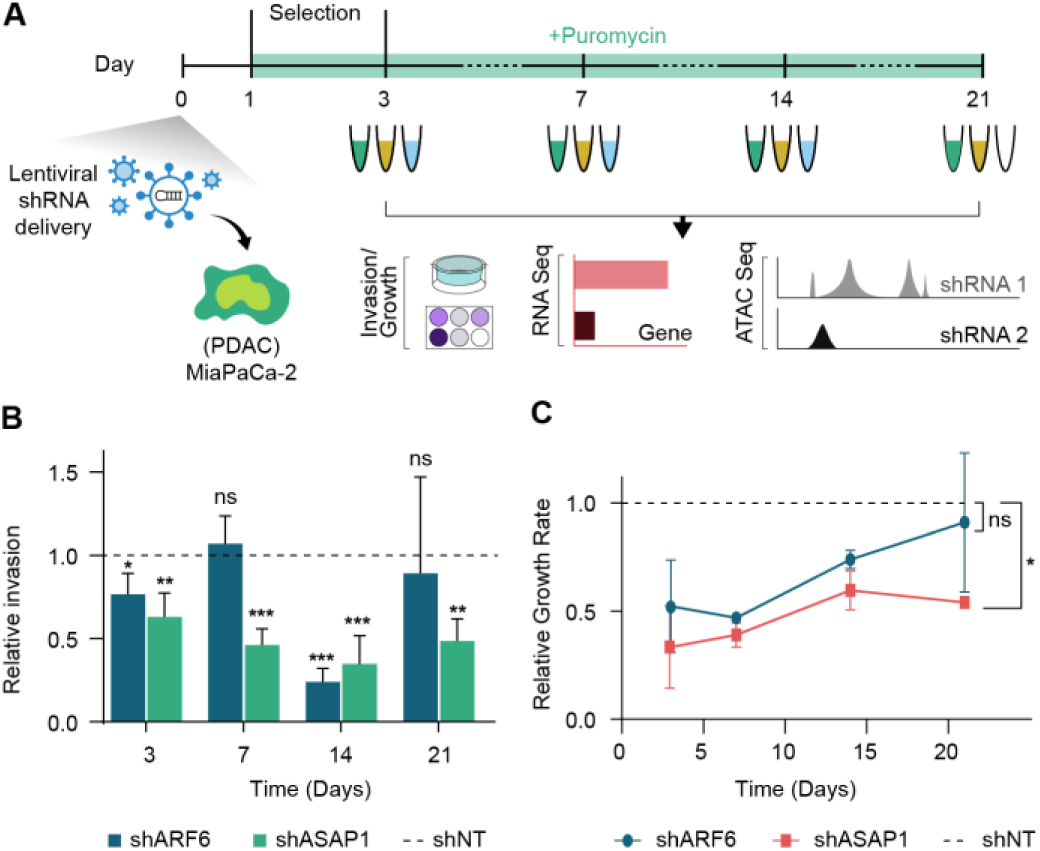
ARF6-depleted MIA PaCa-2 cells adaptively reinstate their invasion and proliferation potential. (A) Schematic of experimental strategy to explore the adaptive effects of downregulating ARF6 or ASAP1 in KRAS-dependent PDAC cells. MIA PaCa-2 cells were treated with shRNAs for ARF6 and ASAP1 and selected using Puromycin for 2 days. (B) Invasion assays were performed using Transwells coated with Matrigel. Assay was performed following treatment with shRNA treatment as in *A*. Invasion was calculated as number of invaded cells at 15 hours normalised to control. (C) Proliferation assay was performed using MTT readout. Assay was performed following shRNA treatment as in *A*. Growth rate was calculated as the normalised cell count on day 2 post seeding compared to day 1 post seeding. Statistics in *B-C* are performed using a T-test (* P<0.05, ** P<0.01, *** P<0.001). Data in *B-C* represents mean ± sd (n=3).

Together, our findings indicates that loss of ASAP1 maintains a lower invasion and proliferation potential, while loss of ARF6 calls for rewiring and adaptive reinstation of invasion and proliferation potential. To harness ARF6 inhibition for the treatment of KRAS-driven cancers it is fundamental to understand which factors allow ARF6 downregulated cells to recover.

### Gene ontology enrichment analysis of time-resolved transcriptomics data confirms the adaptation phenotype of ARF6 downregulated cells

Next, we performed time-resolved RNA sequencing (RNA-seq) for cells treated with shNT, shARF6 or shASAP1 using a similar time-based experimental set up (Figure 1 A). Given the successful, or the lack of, reversion of invasion and proliferation potential in shARF6-, or shASAP1-treated cells, further transcriptomic analysis will reveal important insights into the adaptive processes. A principal component analysis for RNA-seq validated the technical replicates, which were clustered together (Supplementary Figure S2 A).

GO analysis of differentially upregulated genes upon shASAP1 treatment corresponded with expected ASAP1 regulators (Supplementary Figure S2 B). In the early time point on day 3, phosphatidyl inositol-3-phosphate-related, GTPase-related and GEF-related terms were enriched, all of which suggest an upregulation of MAPK and PI3K/AKT pathways likely as positive feedback into downstream ASAP1. On day 7, we identified an enrichment of GOs related to the ER (Supplementary Figure S2 B), to which ASAP1 has previously been associated.^23^ In addition to this, on day 14 and 21, various metabolic processes were enriched (Supplementary Figure S2 B), suggesting possible rewiring attempts to adapt to ASAP1 loss. Among them, various glycoprotein-related terms were enriched which are common regulators of response in perturbed KRAS-downstream signalling in various cancers including PDAC.^24–26^ GO analysis of differentially downregulated genes resulting from shASAP1 treatment, highlighted mostly exclusive GOs for day 3, but more conserved changes through the later time points (Supplementary Figure S2 C). Most downregulated GOs on day 3 reflected processes related to DNA replication, RNA processing, ribosomal RNA related processes and chromosome segregation. These findings agree with our observed reduction in cell proliferation from the MTT-based growth assays (Figure 1 C), and previous literature.^27^ Later time points 7, 14 and 21 highlighted GOs related to GTPase-dependent regulation, transcription-related processes and serine/threonine kinase activity (Supplementary Figure S2 C). Given the KRAS-ARF6-downstream role of ASAP1, and regulation by the MAPK pathway, downregulation of GTPase activity is expected. The downregulation of serine/threonine kinase activity corresponds with a reduction in MAPK signalling which regulates cell proliferation and invasion in cancer cells.^28^ We have shown both these phenotypes to be downregulated in PDAC MIA PaCa-2 cells treated with shASAP1 (Figure 1 B-C). In general, all ontologies downregulated upon shASAP1 treatment, confirm previously established aspects of ASAP1 signalling in cancer and the results of our time resolved in vitro assays.

Enrichment of GOs from differentially regulated genes in cells treated with shARF6 (Figure 2 A-B), also revealed very similar modulations as in shASAP1. Analysis of differentially upregulated genes in cells treated with shARF6 showed an enrichment of deubiquitinase activity, regulation of GTPases and serine/threonine kinase activity at the early day 3 time point (Figure 2 A). As before with ASAP1, the upregulation of genes related to the serine/threonine kinase activity likely establish a positively feedback loop into ARF6 activation. Additionally, ARF6 function is regulated through ubiquitination which further impacts cellular migration and invasion.^29,30^ Importantly on day 7, we observed an enrichment of cell cycle related terms surround the regulation of mitosis and chromosomal segregation (Figure 2 A), which we did not observe upon shASAP1 treatment (Supplementary Figure S2 B). These terms align with our findings that PDAC cells subject to ARF6-depletion, but not ASAP1-depletion, produce an adaptive proliferation response starting at day 7 (Figure 1 C). Later time points, especially day 14 and 21 also highlighted the upregulation of GOs related to vesicular transport between the ER and Golgi (Figure 2 A), functions to with which ARFs have been strongly associated.^31^ Enrichment analysis of differentially downregulated genes in ARF6-depleted cells identified GO terms related to glycoprotein metabolism, ribosome biogenesis, RNA processing and ER-related processes on day 3 (Figure 2 B). As above, KRAS-downstream pathways are heavily regulated by or regulate glycoprotein-related processes.^24,25^ Later time points highlighted the downregulation of various ciliary processes (Figure 2 B). Primary cilia are often regulated by ARF6,^32^ and have been associated with 25% of PDAC patients, all of whom showed higher rates of lymph node metastasis and poor prognosis.^33^

**Figure 2.**
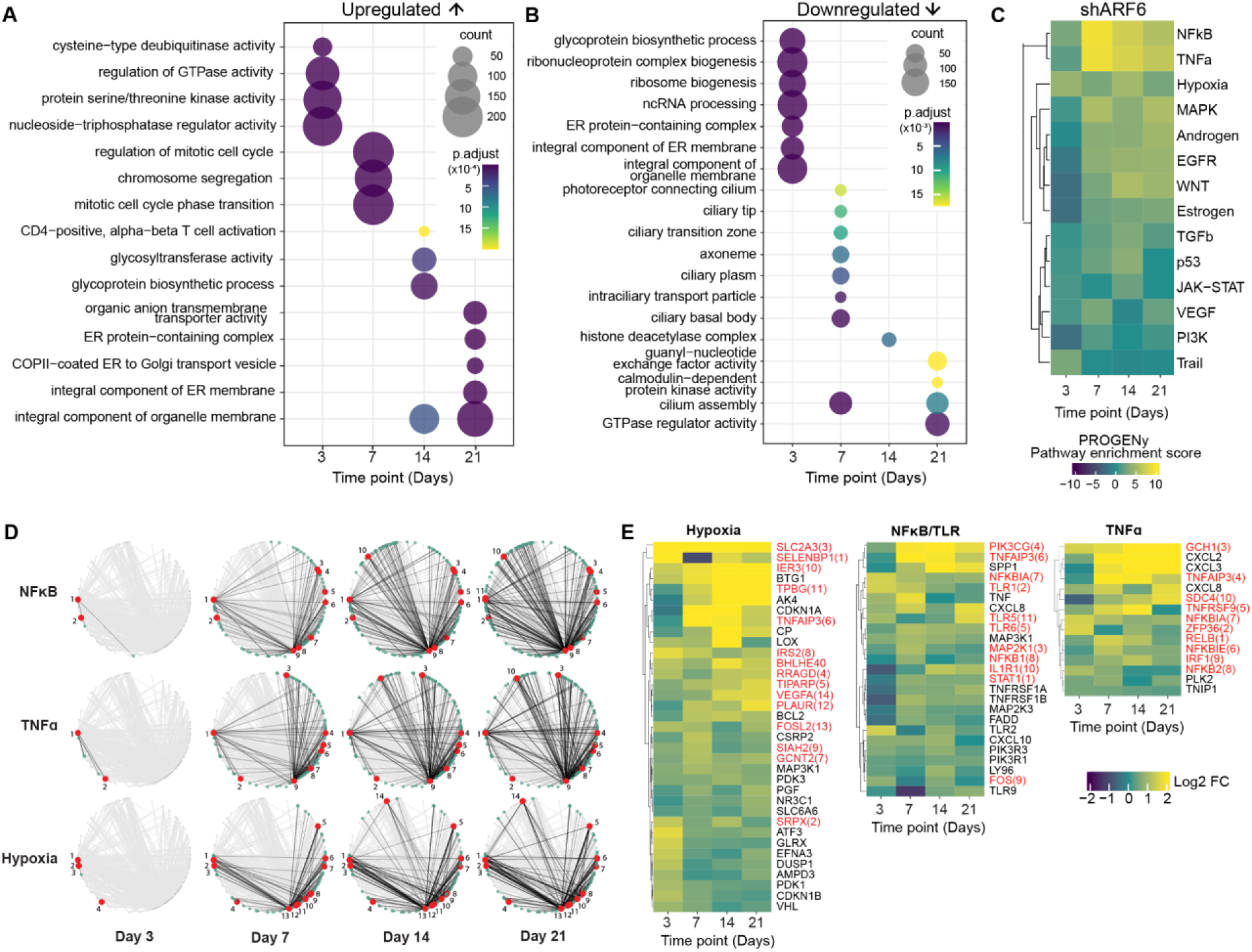
Time-resolved transcriptomic data reveals the upregulation of NFkB, TNFa and Hypoxia pathway in ARF6-depleted MIA PaCa-2 cells. (A) GO enrichment for upregulated genes in shARF6 treatment. (B) GO enrichment for downregulated genes in shARF6 treatment. (C) PROGENy pathway enrichment score from differentially expressed genes in shASAP1-treated cells. (D) PROGENy pathway genes mapped on a PropaNet network created from the enriched TFs and their respective target genes in shARF6-treated cells. Black edges represent all interactions between TFs found in the PROGENy data sets (red nodes) and all their interacting genes (green nodes). See also Supplementary Figure S2 E. (E) Expression of genes of the Hypoxia, NFκB and TNFα pathways from the PROGENy dataset found in our transcriptomic analysis. Red annotations represent TFs found in both the PROGENy and PropaNet datasets as represented in *D*. NFκB gene set was supplemented with genes from the TLR pathways known to feed into NFκB signalling.

Taken together, the GO enrichment analysis of our time-resolved transcriptomics data confirms the adaptation phenotype of ARF6-downregulated cells, which was not observed in ASAP1-downregulated cells.

### Time-dependent transcriptomic analysis shows upregulation of NFkB, TNFa and Hypoxia pathways upon ARF6-depletion

To further explore the transcriptomic data, we employed Pathway RespOnsive GENes for activity inference (PROGENy),^34^ which is a previously published resource that uses publicly available signalling perturbation data to create a dictionary of pathway responsive genes. Overall, when comparing cells treated with shARF6 and shASAP1, we observed large-scale pathway changes only in shARF6 (Figure 2 C, Supplementary Figure S2 D). While there were some pathway perturbations in shASAP1 treatment, they were muted in comparison (Supplementary Figure S2 D). Specifically, in shARF6 treatment, we found major upregulations in the NFκB, TNFα and Hypoxia pathways (Figure 2 C). While these three pathways showed constitutive upregulation following the shARF6 treatment, other pathways such as the Androgen, Estrogen, EGFR and WNT pathways showed an adaptive upregulation (Figure 2 C). Unlike shASAP1 treatment, we did not observe any major downregulation upon shARF6 treatment, which, in general, yielded a much stronger response in terms of upregulation (Figure 2 C, Supplementary Figure S2 D).

Transcription factors (TFs) play a major role in regulating gene expression through enhancer and promoter binding. Hence, changes in gene expression can be directly related to TF activity. To further validate the upregulated PROGENy pathways as a result of shARF6 treatment, we employed PropaNet, a TF network propagating algorithm which is able to consider time-series transcriptomic data and create a minimum interactions network to explain the significant changes.^35^

To identify the TFs that are enriched as a result of ARF6 knockdown, we used ChEA3, a TF enrichment analysis tool.^36^ We obtained a ranked result including 1633 different TFs and their corresponding target genes (supplemental data). From this list, using the ChEA3 TF enrichment score, we shortlisted the top 5% of TFs to generate the PropaNet network. Finally, we mapped specific PROGENy pathway gene sets, which were differentially expressed in our transcriptomic data set, to the PropaNet network. We observed that six pathway gene sets – EGFR, Estrogen, Hypoxia, Androgen, NFκB and TNFα – were represented in the network (Figure 2 D, Supplementary Figure S2 E). More specifically, we found NFκB, TNFα and Hypoxia pathways to be highly enriched within the TF regulatory network (Figure 2 D).

From these pathways, we identified a series of upregulated cytokines (Figure 2 E). These included chemokine family members chemokine (C-X-C motif) ligand (CXCL) 2, CXCL3, CXCL8 and CXCL10. We also identified cytokines interleukin (IL) 1β (IL1B), IL receptor 1 (ILR1) and interferon regulatory factor (IRF) 1, and cytokine-activated factors tumour necrosis factor (TNF) alpha-activated protein (TNFΑIP3) and TNF super family receptor proteins TNFRSF1A and TNFRSF1B. Chemokines and cytokines have been strongly associated with increased cancer cell survival, proliferation and metastasis,^37^ and have been shown to induce angiogenesis (VEGFA, within the Hypoxia pathway, was also found upregulated at later time points in our data)^38^ and cell migration.^39,40^

Together our results from PROGENy pathway enrichment, as well as the PropaNet gene set mapping suggest that the reversion of invasion and proliferation phenotypes in shARF6-treated MIA PaCa-2 cells may be attributed to the upregulation of NFκB, TNFα and Hypoxia pathways.

### Integrating RNA-seq and ATAC-seq reveals TLR-mediated signalling as adaptation mechanisms to ARF6 loss

Transcriptomic analysis reveals that downregulation of ARF6 activates the NFκB, TNFα and Hypoxia pathways as likely adaptations to ARF6 loss. Using the same experimental set up as before (Figure 1 A), we also performed assay for transposase-accessible chromatin with sequencing (ATAC-seq) for shARF6-treated samples collected on days 3, 7, 14 and 21 with the main intention of further streamlining our findings from the transcriptomic data and identifying specific factors which regulate this adaptation. A dimensionality reduction analysis of the ATAC-seq data confirmed agreeable technical replicates (Supplementary Figure S3).

GO enrichment analysis of differentially accessible genes, assigned based on proximity to differentially accessible regions, revealed large scale changes on day 7 (Figure 3 A-B). More specifically, on day 7 we observed a highly represented enrichment of GOs from regions of increased accessibility related to cell cycle (Figure 3 A). These included terms encapsulating chromosome and centromere regions, the mitotic spindle and nuclear division, upregulation of DNA metabolic processes and DNA replication. These terms align with our findings that PDAC cells subject to ARF6-depletion, produce an adaptive proliferation response starting at day 7 (Figure 1 C, Figure 2 A). Time point day 14 highlighted terms related to ribosomes, cell-substrate junctions and focal adhesions (Figure 3 A), all of which play important roles in cytoskeleton-mediated pathways in cancer which regulate migration and invasion,^41,42^ agreeing with our observed recovery of the invasive phenotype in ARF6-depleted MIA PaCa-2 cells (Figure 1 B). Day 21 showed the upregulation of terms related to hypoxia signalling and TF-mediated transcriptional regulation, agreeing with our previous transcriptomic analysis (Figure 2 C-E).

**Figure 3.**
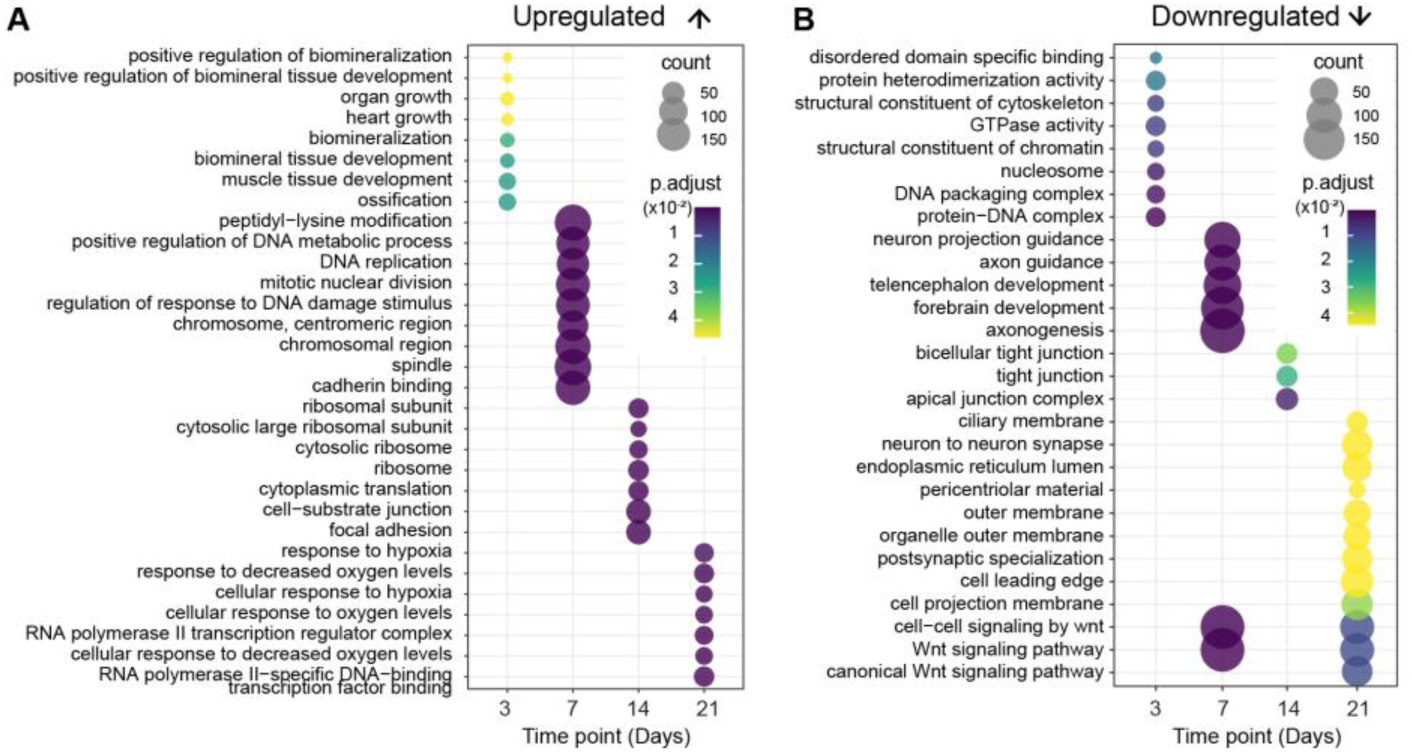
Gene ontology enrichment analysis from ATAC-seq data confirms adaptations in ARF6-depleted MIA PaCa-2 cells. (A) Enrichment of GO terms from differentially higher accessibility regions in shARF6-treated cells in comparison to shNT. (B) Enrichment of GO terms from differentially lower accessibility regions in shARF6-treated cells in comparison to shNT.

Regions of reduced accessibility indicated GO terms specific to ARF6-regulated processes (Figure 3 B). At the early day 3 time point we observed reduced accessibility to fundamental functions of ARF6 such as GTPase activity, cytoskeletal structure and protein regulation (Figure 3 B).^29,31,43^ In addition, we also observed closing of regions related to chromatin structure, DNA packaging and nucleosomes (Figure 3 B), which play a role in cell division,^44,45^ and hence agree with our observed initial reduction in cell proliferation upon ARF6-depletion. Days 7, 14 and 21 highlighted reduced accessibility to various terms related to cytoskeletal and cell junction processes,^15,43,46^ ER-related function^47^ and Wnt signalling,^48^ all of which are ARF6-regulated processes (Figure 3 B).

While various approaches have been previously used to integrate transcriptomic and chromatin accessibility data, few previous works have integrated time-resolved multi-omics data. Recently, work has been done on exploring time-series meta-omics data to understand microbial ecosystems by integrating meta-genomics, meta-transcriptomics and meta-proteomics.^49^ Also, an R-based package, timeOmics, has been recently made available which allows the integration of longitudinal time-course multi-omics data.^50^ Nevertheless, methods that integrate time-dependent multi-omics is still at a very early stage.

To identify specific factors related to adaptations in ARF6-downregulated PDAC, we developed an ad hoc approach to integrate our time-series RNA-seq and ATAC-seq data. ATAC-seq assesses chromatin accessibility, which is an important factor to allow for TF binding to their respective DNA motifs.^51^ Hence, we based our ad hoc integration approach on the enrichment of TFs from differentially expressed genes, and validation of their binding to target gene promoters from chromatin accessibility data. To do this, we used a ChEA3 list of enriched TFs from upregulated genes upon AR6-depletion (Supplementary Figure S4 A) and their respective target genes. We defined the promoter region of each target gene as up to 2kb upstream and 300bp downstream of the transcription start site, and selected for TF-target gene relationships where a TF binding motif was identified within the promoter region. Of note, TFs may have more than one validated binding motif,^52^ however, in our study, to limit the number of hits, we only considered the most highly validated TF binding motifs.

Next, we proceeded to integrate the chromatin accessibility information by comparing the RNA-seq and ATAC-seq datasets and performing a correlation analysis between the expression and promoter accessibility patterns over time. A positive Spearman’s correlation coefficient was considered an agreement between the expression pattern and promoter accessibility for TF binding. From this analysis, we identified 40 TFs (Table 1), and an entire TF-target gene network of over 3000 connections (supplemental data).

**Table 1.** Transcription factors identified from integrating RNA-seq and ATAC-seq. TF and respective target gene count enriched from RNA-seq were validated using ATAC-seq-based MOTIF binding analysis and correlation between expression levels and target gene promoter accessibility. The largest family of TFs and TF subunits that we identified belonged to NFκB pathway and are highlighted in bold.

| TF | Target gene count | TF | Target gene count |
| --- | --- | --- | --- |
| AHR | 211 | ELF3 | 68 |
| MAFB | 176 | **NFKB1** | **68** |
| JUNB | 174 | NR4A2 | 67 |
| BACH1 | 173 | CEBPD | 60 |
| CEBPB | 167 | ETV3 | 59 |
| **NFKB2** | **135** | FOSL1 | 59 |
| BHLHE40 | 134 | PRDM1 | 58 |
| FOSL2 | 124 | ETS2 | 54 |
| **RELB** | **119** | ETV5 | 45 |
| FOXM1 | 104 | BATF3 | 38 |
| E2F1 | 101 | MYBL2 | 38 |
| ATF3 | 100 | FOXD1 | 35 |
| NFE2L2 | 100 | HIF1A | 35 |
| STAT1 | 100 | E2F7 | 26 |
| NFIL3 | 93 | HSF4 | 19 |
| IRF1 | 90 | BHLHE41 | 17 |
| EHF | 89 | NR1H3 | 14 |
| IRF4 | 88 | FOS | 13 |
| KLF5 | 77 | RUNX2 | 12 |
| TP63 | 72 | GATA3 | 11 |

Among these enriched TFs, the largest target gene network was associated with inflammatory signalling pathways. These included the Aryl hydrocarbon Receptor (AHR) which can promote tumour growth through cytokine production such as interleukin-1.^53^ We also identified the Basic Helix-Loop-Helix Family Member E40 (BHLHE40) which can promote colorectal cancer progression through the upregulation of KLF7,^54^ which in turn can regulate NFκB signalling. Indeed, the largest family of TFs and TF subunits that we identified belonged to NFκB pathway – including NFκB1, NFκB2 and RELB (Table 1, bold).

Finally, since we had started with a ChEA3 TF enrichment specifically for the upregulated subset of transcriptomic data, we classified the target genes into constitutively upregulated and adaptively upregulated (Figure 4 A) – and performed Reactome^55^ pathway enrichment (Figure 4 B-C). While adaptively upregulated genes would signify important pathways that are activated in order to maintain or aid the PDAC cell adaptation to ARF6-depletion, constitutively upregulated genes represent the pathways and processes that are critical to initiate the adaptive response to begin with, making them keystones to the PDAC adaptive landscape. In the adaptively upregulated cluster we found genes related to metabolism, cell-ECM interactions, and cell cycle, once again confirming the rewiring process put in place upon ARF6 loss that lead to migration and proliferation recovery (Figure 4 B). The constitutively upregulated genes returned an overwhelming enrichment of TLR and TLR-downstream cascades (Figure 4 C). Of note, TLR2 signalling was enriched repeatedly, in conjunction with TLR4, TLR6 and by itself. We also found MyD88-dependent and –independent signalling patterns, and an activation of the NFκB pathway (Figure 4 C).

**Figure 4.**
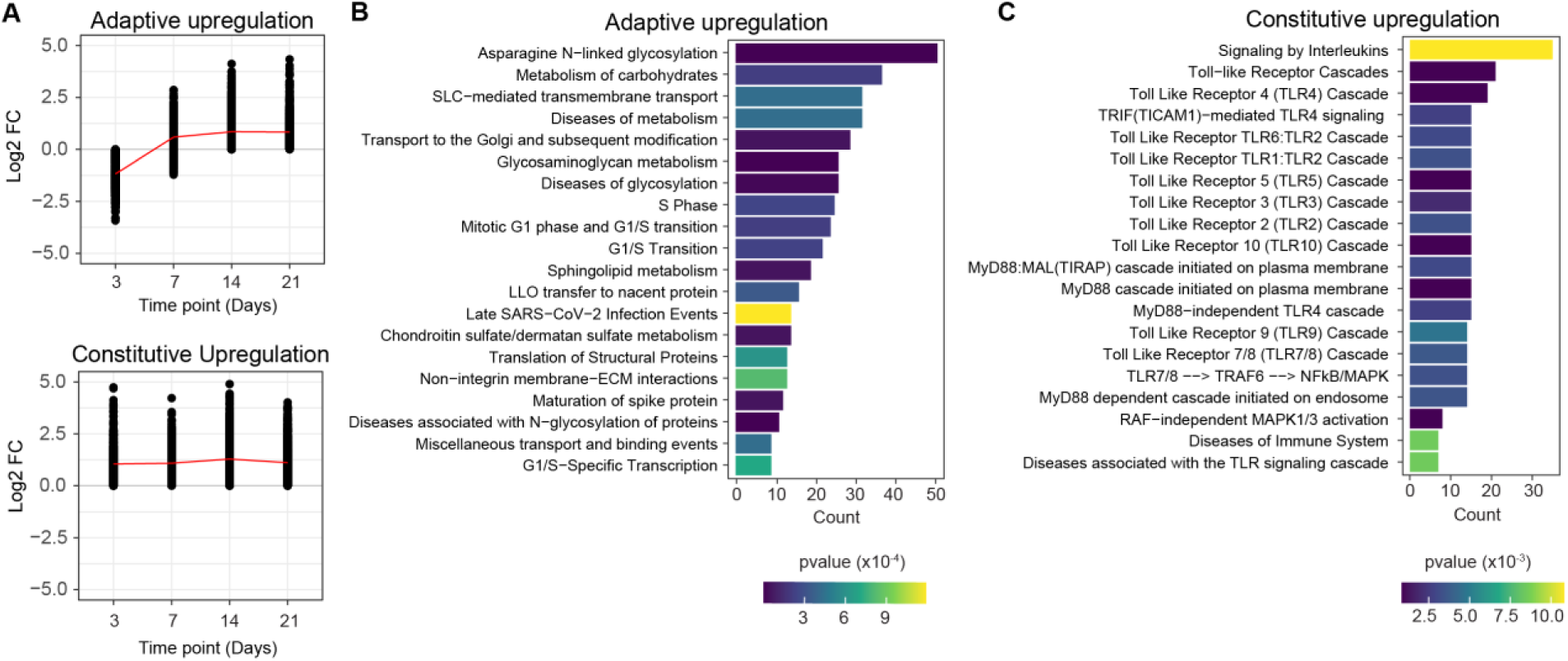
Integration of RNA-seq and ATAC-seq identifies TLR2 as a mediator of adaptation in ARF6-depleted MIA PaCa-2 cells. (A) Upregulated gene clusters from the integration of RNA-seq and ATAC-seq of shARF6 treated samples across time. Adaptively upregulated genes would signify important pathways that are activated in order to maintain or aid the PDAC cell adaptation to ARF6-depletion; constitutively upregulated genes represent the pathways and processes that are critical to initiate the adaptive response, making them keystones to the PDAC adaptive landscape. (B-C) Reactome pathway enrichment of genes from *A* for (B) adaptively upregulated genes, and (C) constitutively upregulated genes.

Overall, from this ad hoc integration of RNA-seq transcriptomic data and ATAC-seq chromatin accessibility data, we identified an overwhelming activation of TLR signalling pathways resulting in NFκB activation. More specifically, we found TLR2 signalling to be highly represented in the pathway enrichment analysis.

### Dual silencing of ARF6 and TLR2 significantly reduces PDAC cell proliferation, spheroid formation and migration in vitro

To evaluate the importance of various TLRs in buffering the loss of ARF6, we performed co-knockdown of ARF6 with TLRs 1-4, TLR9 and MyD88. Our –omics integration revealed TLR2 to be highly represented, and TLR1 and TLR4 have been studied to signal through similar and complementary pathways. We also tested TLR3 and TLR9, both of which are known to signal from endosomal compartments, but using distinct downstream cascades. Finally, we also included MyD88, which is a major TLR-downstream adaptor molecule.

To design the in vitro experiments, we revisited our preliminary Matrigel-invasion and proliferation assays together with the results of the time-resolved OMICS integration data. MIA PaCa-2 cells were transduced with shARF6 and shTLR lentiviruses either in combination or by themselves, and selected for 7 days using antibiotics, when MIA PaCa-2 cells showed signs of recovering their proliferation and invasion phenotypes (Supplementary Figure S5 A, Figure 1 B-C). Quantitative real-time PCR confirmed the successful downregulation of ARF6, TLRs and MyD88 expression levels in comparison to control cells (Supplementary Figure S5 B).

Cell proliferation assays were based on readings from two consecutive days, where the value from the second day was normalised against the value from the first day to measure the proliferation rate over a period of 24 hours. Typically, MIA PaCa-2 cells have a doubling time of approximately 18 hours. In agreement with this, we detected a proliferation rate of 2.8x for wild-type cells (Figure 5 A). Upon treatment with shARF6, the average proliferation rate was significantly decreased to 2.0x (Figure 5 A). Downregulation of TLR2 and MyD88 by themselves also showed significant reduction of mean cell proliferation in comparison to control cells (Figure 5 A). TLR2 treatment reduced the rate to 1.9x, and MyD88 treatment showed similar results as shARF6 treatment (Figure 5 A). While some other TLRs such as TLR3 and TLR4 also exhibited a reduction in proliferation rates to 1.7x and 1.9x respectively, we observed high variability between biological replicates. Co-knockdown together with ARF6 identified various TLRs showing significantly reduced proliferation rates compared to WT (Figure 5 A). However, only TLR2 and ARF6 co-knockdown showed not only a reduction in proliferation (1.2x) in comparison to the WT control (2.8x), but also a combined effect relative to the ARF6 (2.0x) and TLR2 (1.9x) treatments by themselves (Figure 5 A).

**Figure 5.**
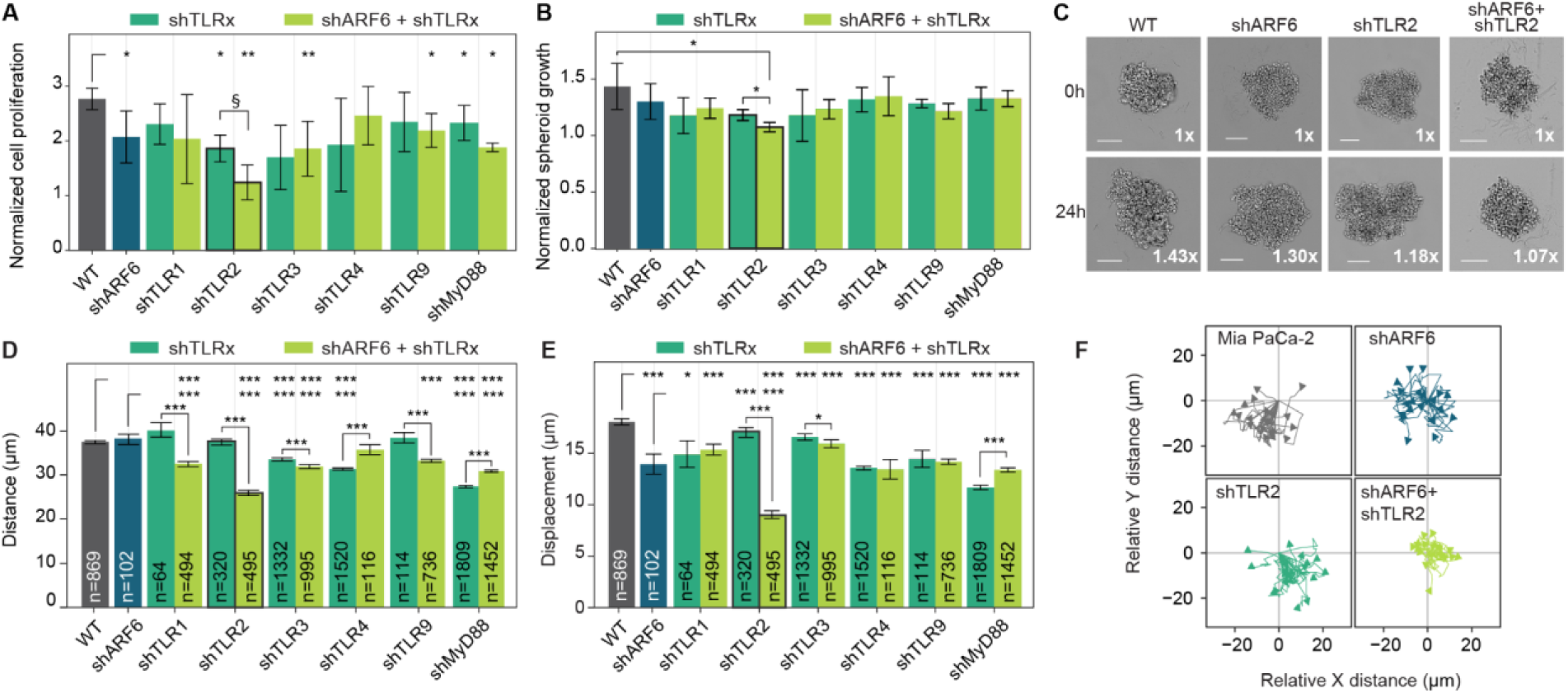
ARF6 and TLR2 co-knockdown inhibits MIA PaCa-2 cell proliferation and migration in vitro. (A) Quantification of cell proliferation using the crystal violet assay following shRNA treatment for either single infections of ARF6 or TLRs, or co-infection of ARF6 and TLRs. Graph represents mean ± sd (n=3). (B) Quantification of relative spheroid growth using shRNAs for ARF6 or TLRs, or the co-infection of ARF6 and TLRs. Graph represents mean ± sd (n=3). (C) Representative images of spheroids treated with ARF6, TLR2 or, ARF6 and TLR2. (D-E) High-throughput quantification of (D) total distance travelled or (E) total displacement by individually tracked cells from their point of origin over 10 hours, shown as mean ± sd. (F) Representative migration trajectories of 20 sampled cells per condition. Statistics in *A, B, D* and *E* were performed using the Wilcoxon test (§ P=0.058, * P<0.05, ** P<0.01, ** P<0.001, *** P<0.001). Unless indicated, comparisons are not significant.

Next, to understand proliferation in a more physiologically relevant context, we used 3D spheroid growth assays. As with the proliferation assays, two measurements were taken on consecutive days and the spheroid area was normalised against the value from the first day to obtain the proliferation rate over 24 hours. MIA PaCa-2 WT cells increased spheroid area by 1.43x on average (Figure 5 B-C). This increase was limited to an average of 1.3x and 1.18x for cells treated with shARF6 and shTLR2 respectively (Figure 5 B-C). Of note, the combinatorial treatment of shARF6 together with shTLR2 resulted in a significantly stinted spheroid growth limiting the proliferation to only 1.07x, which was also significantly lower than the single ARF6 or TLR2 knockdowns (Figure 5 B-C). No other shTLR treatment, by themselves or in combination with ARF6 downregulation resulted in a significantly reduced spheroid growth (Figure 5 B).

Finally, given the importance of the KRAS-ARF6 in enabling migration, we investigated the impact of downregulating ARF6 and TLR expression levels in MIA PaCa-2 cells on cellular mobility. To do this we performed high-throughput 2D cell migration assays using time-lapse imaging. MIA PaCa-2 cells treated with shRNA for ARF6, TLRs or both, were individually tracked for a period of up to 10 hours and the average distance and displacement were compared. MIA PaCa-2 wild-type cells migrated an average distance of 37.4 µm (Figure 5 D), and showed an average displacement of 18.1 µm (Figure 5 E). Upon treatment with shARF6 the average distance remained similar at 38.7 µm (Figure 5 D), while the average displacement was significantly reduced to 0.77-fold or 14 µm (Figure 5 E), suggesting that shARF6 leads to cells moving about their point of origin, rather than migrating further away, in comparison to WT cells. Individual knockdown of some TLRs also reduced both distance and displacement significantly in comparison to WT (Figure 5 D-E). However, in most cases either we did not observe any synergistic or additive effect of a co-knockdown of ARF6 and the TLR; or, we did not observe consistent effects on distance and displacement. Most strikingly, co-knockdown of TLR2 with ARF6 reduced the average migration distance to 0.7-fold (Figure 5 D, F), and average displacement to 0.5-fold (Figure 5 E-F). These distance and displacement values were also significantly lower when compared to their respective shARF6 or shTLR2 individual controls.

Finally, we looked at the ARF6 and TLR2 co-knockdown in terms of a distance-displacement multivariate analysis at a single-cell resolution (Supplementary Figure S5 C). Wild-type MIA PaCa-2 cells exhibited large migration distances resulting in a large displacement (Supplementary Figure S5 C). Compared to this, both shTLR2 and shARF6-treated cells exhibited significantly reduced migration compared to the wild-type control, but only mildly different from one another (Supplementary Figure S5 C). In stark contrast to all three of the above treatments, the co-knockdown of the ARF6 and TLR2 significantly impaired migration on both axes for the majority of the population (Supplementary Figure S5 C).

In conclusion, our results from the in vitro assays for 2D cell proliferation, 3D spheroid growth and 2D cell migration, all validate TLR2 as an important factor compensating for the loss of ARF6 signalling.

### Dual silencing of ARF6 and TLR2 leads to a reduction in tumour growth in vivo

TLR2 silencing has a strong impact on ARF6-depleted pancreatic cancer cells in terms of in vitro 2D cell proliferation and 3D spheroid growth. To evaluate if these effects translate to in vivo systems, we used a tumour xenograft mouse model. Luciferase+ FACS sorted MIA PaCa-2 cells (Supplementary Figure S6 A) were treated with shRNA for ARF6, TLR2 or both, for a period of 7 days. Following this, we performed subcutaneous injections in equal numbers of male and female athymic immunodeficient mice. Visible and quantifiable tumours appeared in the shNT-injected mice approximately 2-weeks post injection, while shARF6, shTLR2 or shARF6+shTLR2-injected mice only showed the first signs of tumour at approximately 3-weeks post injection. Of the 12 possible tumours, 6 in females and 6 in males, shNT produced tumours in all 12 cases (Figure 6 A-B) and tumour volume increased at an average rate of 13.7% per day (Figure 6 C). As a result, shNT tumours reached an average volume of 550 mm^3^ and a maximum tumour volume of 1000 mm^3^ in a span of 7-weeks post-injection (Figure 6 C), at which point the experiment was concluded. In contrast to control conditions, shARF6 or shTLR2 treatment alone resulted in 6 out of 10, and 7 out of 12 tumours respectively (Figure 6 A-B). Both these conditions showed significantly slower average growth rates of 9.4% and 8.8% per day respectively, in comparison to shNT treatment (Figure 6 C). Additionally, while the downregulation of ARF6 and TLR2 alone strongly affected tumour growth in vivo, their combinations almost abrogated tumour proliferation, showing only a single tumour with a final volume of 19.8 mm^3^, which was significantly lower than the average tumour volume of the other three conditions (Figure 6 C). Luciferase imaging prior to tumour extraction also confirmed our results (Supplementary Figure S6 B).

**Figure 6.**
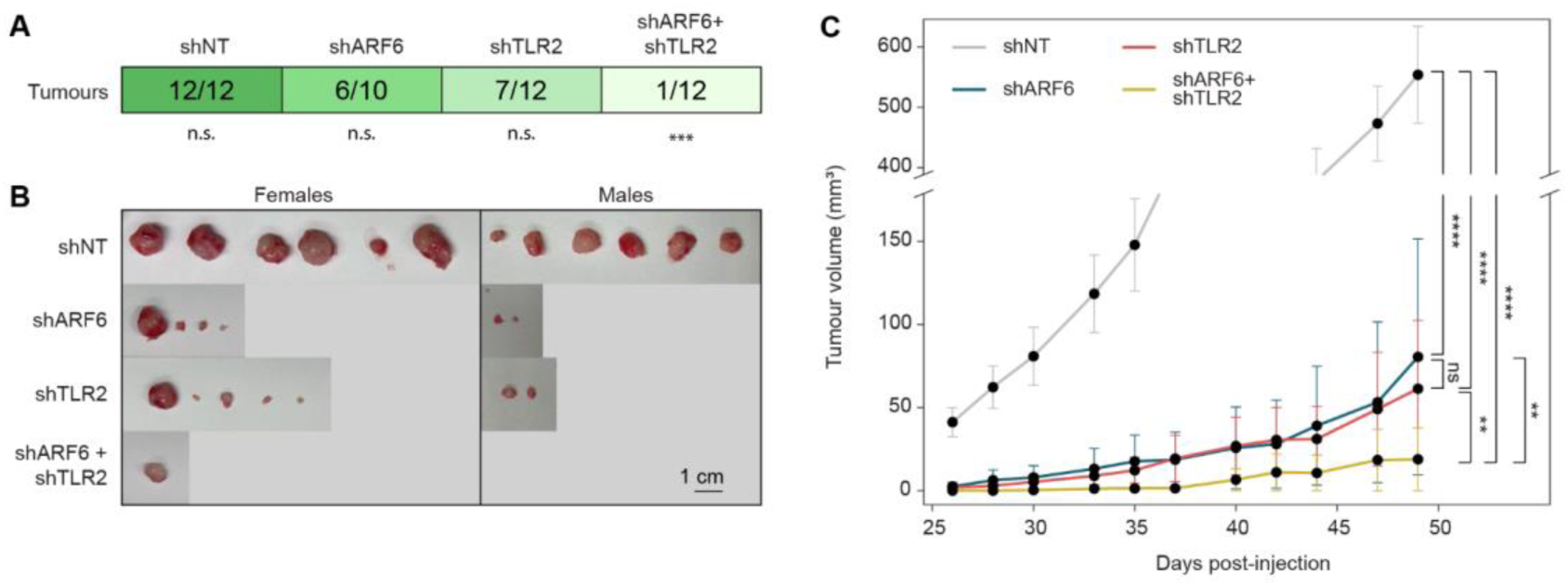
ARF6 and TLR2 co-knockdown inhibits MIA PaCa-2 tumour growth in vivo. (A) Table showing quantification of total tumours observed as a fraction out of the total possible tumours per condition. Experiment was performed in 3 male and 3 female mice, where 1×10^6^ treated MIA PaCa-2 cells were subcutaneously injected in both left and right flanks and tumours monitored over time (total possible tumours n=12). One animal died during the experiment in ARF6 treatment (total possible tumours n=10). (B) Pictures of the tumours (n=10-12 tumours per condition) extracted at the end of the experiment from mice. (C) Tumour volume (n=10-12 per condition) over time. Data are shown as mean ± sd. Statistics in *A* was performed using Chi-squared test to compare expected and observed tumours in each condition. (* P<0.05, ** P<0.01, *** P<0.001). Statistics in *C* was performed to evaluate growth trend over time using the Mann-Whitney-Wilcoxon test (* P<0.05, ** P<0.01, *** P<0.001).

In conclusion, our mouse xenograft models show that both ARF6 and TLR2 play important roles in tumour growth in vivo. While the loss of either of them individually causes slower tumour growth rate, their simultaneous knockdown almost completely abolishes tumour growth.

### Dual silencing of ARF6 and TLR2 leads to a reduction in metastatic potential in vivo

Pancreatic cancer cells are well characterised to be aggressive and invasive. To better understand the reliance of ARF6-depleted cells on TLR2 in the context of metastasis, we employed a zebrafish xenograft model (Figure 7 A).^56^ GFP+ FACS sorted MIA PaCa-2 cells (Supplementary Figure S7 A) treated with shRNA for 7 days as before, were injected into zebrafish larvae at 2 days post fertilisation (2-dpf). Transgenic animals (carrying the cassette fli:GAL4; UAS:RFP) were used to visualise the circulatory system into which the cells were delivered by injections at the duct of Cuvier (Supplementary Figure S7 B). Live imaging of larvae was then performed on days 7 and 8. Day 8 readings were normalised against day 7 readings to assess the relative metastatic potential of MIA PaCa-2 cells.

**Figure 7.**
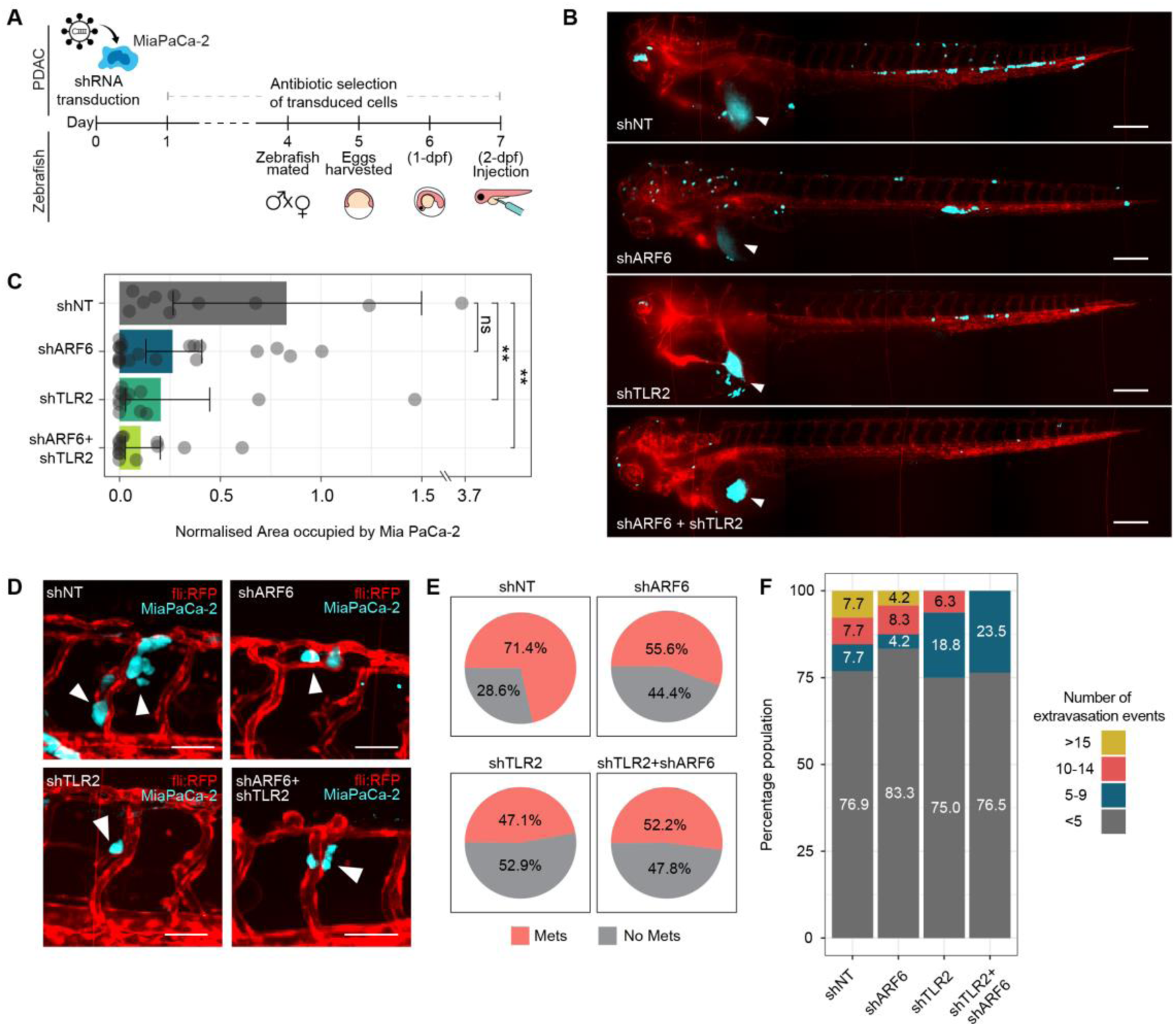
ARF6 and TLR2 co-knockdown deters MIA PaCa-2 metastatic potential in vivo. (A) Schematic representation of in vivo zebrafish xenograft experimental timeline highlighting PDAC cell treatment, and simultaneously the development of zebrafish embryos as required for injections. PDAC cells were treated with shRNAs for 7 days including antibiotic selection, prior to injection. Zebrafish were mated accordingly to obtain 2-days-post-fertilisation (2-dpf) embryos on day 7. (B) Representative images 24 hours post-injection of transgenic (fli:GAL4; UAS:RFP) zebrafish injected with GFP+ MIA PaCa-2 (pCMV-KOZAK-EGFP) cells treated with shRNA for NT control, ARF6, TLR2 or both in combination. White arrows represent the site of injection at the duct of Cuvier. MIA PaCa-2 cells are in blue, and zebrafish vasculature is marked in red. See also Supplementary Figure S7. (C) Quantification of the area occupied by the MIA PaCa-2 cells 24 hours post-injection normalised to 0 hours post-injection. Cells still resident within the site of injection were excluded during analysis. (D) Images of shRNA-treated MIA PaCa-2 cells extravasating (white arrows) out of zebrafish vasculature (red). (E) Proportion of the zebrafish population showing at least 1 extravasation event (Mets) or no extravasation events (No Mets) (F) Quantification of the extravasation events per animal. Number of extravasation events per animal were binned into four categories <5, 5-9, 10-14 and >14. Scale bars in *B* represent 200 µm. Scale bars in *F* represent 50 µm. Statistics in *C* were performed using the Wilcoxon test (* P<0.05, ** P<0.01)

In comparison to the number of circulating cancer cells immediately post-injection, approximately 83% of the population were still present in circulation 24 hours later (Figure 7 B-C). In contrast to this, only 26% of shARF6-treated cells and 20% of shTLR2-treated cells remained in circulation 24 hours-post injection (Figure 7 B-C). Finally, the simultaneous downregulation of ARF6 and TLR2 showed the strongest reduction in cell survivability with only 10% cancer cells remaining in circulation (Figure 7 B-C).

We therefore quantified the extravasation potential of treated MIA PaCa-2 cells by calculating the number of locations per animal where the cancer cells had metastasised into surrounding tissue (Figure 7 D). We found that in control conditions, greater than 70% of animals developed extravasation events (Figure 7 E). The other three treatments - shARF6, shTLR2 and their combination - all showed a similar reduction in extravasation events, with approximately 50% of animals positive for metastasis (Figure 7 E). We grouped these events into high metastatic (>15 extravasations), medium-high metastatic (between 10 and 14 extravasation events), medium-low metastatic (between 5 and 9 extravasation events) and low metastatic (<5 extravasation events). We observed that all conditions had a comparable number of low metastatic cases, which was the most representative phenotype (Figure 7 F). However, when comparing higher metastatic potentials, we observed that the combinatorial downregulation of ARF6 and TLR2 eliminated the presence of either medium-high or high metastatic cases (Figure 7 F).

Overall, the zebrafish metastasis experiments demonstrated that downregulation of ARF6 and TLR2 in combination reduced cell survivability and invasive potential, which was consistent with the in vitro observations.

## DISCUSSION

KRAS-dependent PDAC is one of the most aggressive cancers with poor prognosis due to late stage detection. Greater than 90% of PDACs are diagnosed past Stage I, making them exceedingly difficult to treat medically. The advantage of a mutant form of KRAS driving PDAC is that it is targetable using mutation-specific drugs and small molecules without eliciting a systemic response. AMG510 is a recently FDA-approved drug that is being tested in clinics as potential KRAS G12C therapeutic strategy.^4,57^ Nevertheless, recent work has shed light on the mutagenic side effects of AMG510 treatment.^5^ Targeting major master regulators such as KRAS in malignant cells leads to acquisition of additional mutations in not only RAS but also other important pathways which are capable of resistance and malignancy. Therefore, identifying an alternate and effective strategy to target advanced KRAS-dependent PDAC is urgently warranted.

### ARF6, but not ASAP1 downregulation leads to PDAC adaptation

A strong approach is the targeting of key signalling cascades downstream of KRAS, which can inhibit important PDAC aggressiveness-enabling processes. One such axis is given by ARF6-ASAP1 signalling. ARF6 is a small GTPase of the Ras family of GTPAses and is a major mediator of KRAS signalling. Loss of ARF6 has been studied to impair proliferation, migration and invasion,^13–19^ which in turn has prompted the development of ARF6 inhibitor NAV-2729,^20^ and is considered as a potential therapeutic target.^21^ For instance, ARF6 is an inhibitor of the RLS3-induced ferroptosis and a promoter of gemcitabine resistance in PDAC.^17^ Additionally, ARF6 function may also act as a positive feedback loop for ERK signalling, which is fundamental in various MAPK pathway processes in cancer.^58^ At phenotypic level, ARF6 also regulates various cytoskeletal processes which eventually promote cancer cell migration, and cell invasion – both critical processes for metastasis.^13–15^

Indeed, we also observed that the shRNA-mediated downregulation of ARF6 resulted in stinted proliferation and Matrigel invasion in vitro. Importantly, our study, unlike most previous studies, introduces a temporal dimension. We showed that the loss of ARF6 in KRAS-addicted PDAC leads to an adaptive response, which eventually reverts the proliferative and invasive potential to control levels. In stark contrast to the perturbation of ARF6, the loss of ASAP1, which is a KRAS-endpoint and ARF6-downstream effector, did not result in a reversion of proliferative nor the invasive potential over time, suggesting that perturbations applied at different hierarchical positions within the same KRAS-downstream axis have differential responses.

### TLR signalling as an adaptation to ARF6 loss

TLRs have been associated with both anti-tumorigenic effects, but also pro-tumorigenic effects. Through the ad-hoc integration of RNA-seq and ATAC-seq results of shARF6 samples, we identified a series of TLR-related processes to be upregulated across all the time points tested – day 3, day 7, day 14 and day 21. Primarily, we observed the TLR2 cascade to be most prominently represented, albeit with common genes shared with various other TLR cascades.

An increase in TLR expression does not necessarily imply TLR signalling, and similarly an enrichment of the TLR signalling pathway does not necessarily imply a transcriptional upregulation of the respective TLR. Indeed validation experiments to obtain synergistic effects between co-knockdown of TLRs and ARF6, where we tested multiple TLRs, only validated TLR2. TLR2 is a unique TLR, in that it forms both homodimers^59^ and also heterodimers with more than one type of TLR.^60,61^ In that context, we tested TLR1 and TLR4 downregulation as well, but did not see any significant effects, suggesting that it is likely a TLR2 homodimer-mediated signalling.

TLR2-downstream cascades have been identified as major pro-tumour signalling pathways. In response to radiotherapy, TLR2 signalling stimulated by High mobility group box 1 protein (HMGB1) from dying cells, has been shown to accelerate metastasis in pancreatic cancer, through EMT and upregulation of the PI3K/AKT pathway.^62^ TLR2-MyD88 signalling has also been associated with poor breast and colon cancer prognosis through cell-intrinsic pathways.^63^ In hepatocellular carcinoma TLR2 upregulation activates a MyD88-dependent NFκB pathway which promotes invasion.^64^ Similarly, in most cases, TLR2 signals via the canonical MyD88-mediated pathway. Interestingly however, in our attempt to validate MyD88 we only observed a synergistic effect together with ARF6 knockdown in 2D proliferation assays. In 3D spheroid growth assays we did not observe any significant differences. And, in migration assays we observed an inverted trend where MyD88 knockdown alone reduced cell migration more than the reduction by a co-knockdown with ARF6. Overall, this suggests that TLR2 in MIA PaCa-2 PDAC cells, upon loss of ARF6, likely enables adaptation through a MyD88-independent pathway.

MyD88-independent signalling has so far been identified for various TLRs. TLR4 signalling via a MyD88-independent pathway has been shown to activate the NFκB and interferon pathways.^65^ In fact, ARF6 has been found to play an important role in regulating TLR4 signalling through MyD88 and also via the MyD88-independent pathway.^66^ In contrast, ARF6 plays no role in TLR2 signalling.^66^ Both TLR3 and TLR4 are also well established to signal through a TRIF/TRAM-dependent manner to activate downstream processes.^67,68^ While TLR3 can function with TRIF alone, TLR4 requires both TRIF/TRAM mediation. Indeed, in our Reactome pathway enrichment of constitutively upregulated genes, we found terms related to TLR-signalling through a MyD88-independent pathway. In general, a ubiquitination-dependent regulation is important for balancing MyD88-dependent and –independent pathways which signal through TRIF (conditionally TRIF3 or TRIF6), TRAMs and TAK.^30^ While it is not common for TLR2 to signal in a MyD88-independent pathway, within the scope of this study, we found little evidence to show that TLR2 compensation for ARF6 loss was being mediated by MyD88. Only one study has identified that during infection by *Porphyromonas gingivalis*, a gram negative bacteria, TLR2 can upregulate TNFα and cytokines in a MyD88-independent manner.^69^ Together, our study suggests a new axis given by MyD88-independent, TLR2 signalling in KRAS-dependent ARF6-depleted PDAC.

### TLR2-activated NFκB, TNFα and Hypoxia pathways promote tumour growth and metastasis

The pro-tumorigenic activation of TLR2 signalling proceeds to activate NFκB, TNFα and other pathways.^70^ In agreement, using PROGENy pathway enrichment we found the NFκB, TNFα and Hypoxia pathways to be maximally enriched across all the time points upon ARF6 depletion. In colorectal carcinomas, TLR2 stimulation results in autocrine signalling of TNFα, which induces pro-cancer signalling.^71^ In gastric cancer, TLR2 activation can lead to an increased signalling across the NFκB, PI3K/AKT and ERK-activated MAPK pathways, leading to accelerated tumorigenesis.^72^ TLR2 can also promote cell migration, invasion and colony formation in A549 and H1299 human lung cancer cells via NFκB, by upregulating chemokines, cytokines, MMPs and TNFα.^73,74^ Moreover, TLR2-dependent NFκB can then further continue to signal downstream and upregulate Hypoxia pathway regulator Hypoxia Inducible Factor 1 Subunit Alpha (HIF1α).^75^ HIF1α was also found from our data set upon TF enrichment analysis. Moreover, HIF1α and TLR2 have been specifically associated with malignant transformation in early PanINs,^76^ together supporting a TLR2 dependent maintenance of tumorigenicity.

In our study, when looking into the differentially expressed genes which led to the PROGENy enrichment for NFκB, TNFα and Hypoxia pathways, and also mapped onto the PropaNet-based TF regulatory network, we identified several chemokines and cytokines which are critical in the tumour microenvironment for cellular crosstalk and signalling.^77^ These included IL1R1, CXCL2, CXCL3, CXCL8, CXCL10, and IRF1. We also found TNF, TNFRSF1, TNFRSF2 and TNFΑIP3 members of the TNFα pathway that use feedback loops to cross-regulate NFκB signalling^70^ and eventually suppress anti-tumour immune responses.^78^ We also identified the upregulation of PI3K/AKT-downstream PI3K catalytic subunit gamma (PIK3CG), which is an important mediator for the regulation of NFκB signalling in cancer.^79^ Together this supports the hypothesis that TLR2 activation likely leads to the downstream upregulation of the NFκB, TNFα and Hypoxia pathways.

However, within the scope of this study, we have not assessed whether TLR2 activation leads to NFκB, TNFα and Hypoxia pathway activation, or whether the activation of the pathways leads to a TLR2 signalling and as a result a positive feedback loop. Although literature generally points towards the former as the more likely in the context of pro-tumorigenic signalling, future explorations can shed light on the exact mechanisms. TNFα for instance may signal in an autocrine manner to activate TLRs,^80^ or may act on TLR-downstream factors to activate them.^70,81,82^

### Co-targeting of ARF6 and TLR2 as a novel therapeutic strategy in KRAS-dependent PDAC

Various independent studies have highlighted the role of ARF6 in drug resistance and as a result, targeting of ARF6 is now a seriously considered therapeutic strategy for breast, gastric, renal and pancreatic malignancies (reviewed by Dejuan Sun et al.).^21^ ARF6 with ZEB1 has been shown to confer drug resistance in breast cancer against Gemcitabine, 5-fluorouracil and Temsirolimus.^46^ In renal cell carcinomas it has been linked with resistance against 5-fluorouracil.^83^ And in PDAC, tested using MIA PaCa-2 cells, it has been linked with Gemcitabine resistance.^17^ Silencing or knock out of ARF6 in these studies re-sensitised cancer cells to their respective drug treatments.

In our study, we have shown that the silencing of ARF6 alone may not be an ideal strategy because of the acquired time-dependent adaptation mechanisms that lead to the reversion of proliferation and invasion potential. In fact, various current therapeutic approaches have acknowledged the need for dual targeting strategies. For PDAC patients, one such combination of drugs in phase Ib clinical trials is the CD40 agonist monoclonal body APX005M together with gemcitabine and nab-paclitaxel.^84^ In another study, panitumumab, an anti-EGFR drug and trastuzumabm, an anti-HER2 therapy significantly enhanced the effect of trametinib, a MEK1/2 inhibitor in deterring proliferation in patient-derived xenograft models of PDAC.^85^ Similarly, we found that the dual downregulation of ARF6 and TLR2 leads to an augmented deterrence of tumour proliferation in MIA PaCa-2 xenograft models in mice. Not only did we find that the co-targeting of ARF6 and TLR2 reduced tumour proliferation rates, but it practically abolished tumour growth altogether, with only 1 out of 12 animals developing a tumour. In addition to that, our zebrafish model of metastasis also identified strong trends towards a reduced metastasis potential in MIA PaCa-2 cells with the dual inhibition in comparison to controls, impacting various aspects of metastasis including survival in circulation and extravasation.

Together, our findings concretely suggest the dual knockdown of ARF6 and TLR2 as a new therapeutic strategy in the context of KRAS-dependent cancers, and specifically PDAC. Nevertheless, further studies using commercially available ARF6 inhibitor NAV-2729^20^ and TLR2 inhibitor C29^86^ will be vital to validate our findings.

## Conclusion

KRAS is mutated in over 90% of pancreatic cancers leading to the initiation and progression of PDACs.^11,87^ One therapeutic strategy is to target key KRAS-downstream signalling axes. One such axis is given by ARF6-ASAP1 signalling, which plays an important role in PDAC proliferation, migration, invasion and immune evasion.^15^ In this study, we show that KRAS-dependent PDAC, subject to the downregulation of ARF6 or ASAP1, which hierarchically represent an intermediary and a downstream effector of KRAS signalling respectively, show different cellular responses. We show that only ARF6-depletion, but not ASAP1-depletion, elicits an adaptive response from PDAC cells. This adaptive response is mediated by TLR2 signalling and leads to the upregulation of NFκB, TNFα and Hypoxia signalling pathways. In vitro validation experiments using 2D proliferation assays, 3D spheroid growth assays, and 2D cell migration assays all showed inhibited phenotypes upon co-knockdown of TLR2 and ARF6. Importantly, our in vitro assays did not conclude a need for MyD88, which is a TLR2 signalling mediator, hence suggesting a MyD88-independent signalling response, which remains to be investigated.

ARF6 has been considered as a potential therapeutic target due to its multifaceted role in cancer progression.^21^ We show here that targeting ARF6 alone may not be sufficient, leading to acquired tumour resistance. Co-knockdown of TLR2 with ARF6 showed profound effects in vivo, almost abolishing tumour growth in a mouse xenograft model. Finally, using a second in vivo xenograft model in zebrafish, we showed that the co-knockdown of TLR2 and ARF6 also have the ability to decrease PDAC metastatic potential. Through this study, we identify TLR2 as a major regulator of mechanistic adaptation in ARF6-depeleted PDAC cells and show that the co-targeting of these two factors may offer a novel therapeutic approach.

## ACKNOWLEDGMENTS

We would like to thank the CRG Genomics Unit for their support with sequencing experiments and the CRG/UPF Flow Cytometry Unit for their support with FACS experiments. We thank all the Sdelci lab members for their discussions and critical feedback on this manuscript.

## AUTHOR CONTRIBUTIONS

Conceptualization: RG, SS

Methodology: RG, SS

Software: RG, UG, DD, NPL

Validation: RG, SD, JUS, FP

Formal analysis: RG, SD, DD, UG, NPL

Resources: RG, DD, UG, NPL

Investigation: RG, SD, JUS, FP, AGZ, LE, LG, CC

Data curation: RG, DD, UG, NPL

Writing - original draft: RG, SS

Writing - review & editing: RG, SS, FP, JUS, AJ

Visualisation: RG

Supervision: SS

Project administration: RG, SS

Funding acquisition: SS, VR

## METHODS

### Cell culture

Unless otherwise mentioned, MIA PaCa-2 (ATCC), Panc-1 (ATCC) and HCT-116 (ATCC) cells were maintained in complete Dulbecco’s Modified Eagle Medium (DMEM, Gibco) containing 10% Fetal Bovine Serum (FBS, Gibco) at 37°C and 5% CO2. Cell washes were performed with phosphate buffered saline (PBS) and detachment was done using Trypsin (Gibco). All the cell lines were tested negative for mycoplasma contamination several times during the study.

### Data availability

All data relevant for the work produced in this manuscript has been made available through the supplemental data file, on the GEO repository (GSE247513), or at https://github.com/SdelciLab/arfAdapt.

### shRNA vectors and sequences

shRNA-mediated knockdown was performed using Sigma MISSION shRNA constructs for ASAP1 (TRCN0000123118) and ARF6 (TRCN0000048003). Bacterial stubs were cultured and maxi-prepped using the QIAGEN Plasmid Plus Maxi Kit (Qiagen 12963) and as per manufacturer’s directions. For all other shRNAs used in this study (Table 2), following Addgene protocol, oligonucleotides were ordered (Integrated DNA Technologies) and cloned into the pLKO.1 (addgene #8453) backbone containing a Puromycin resistance gene. For controls, either untreated wild-type or Sigma MISSION non-targeting shNT vector (Sigma-Aldrich Shc002) was used.

**Table 2.**
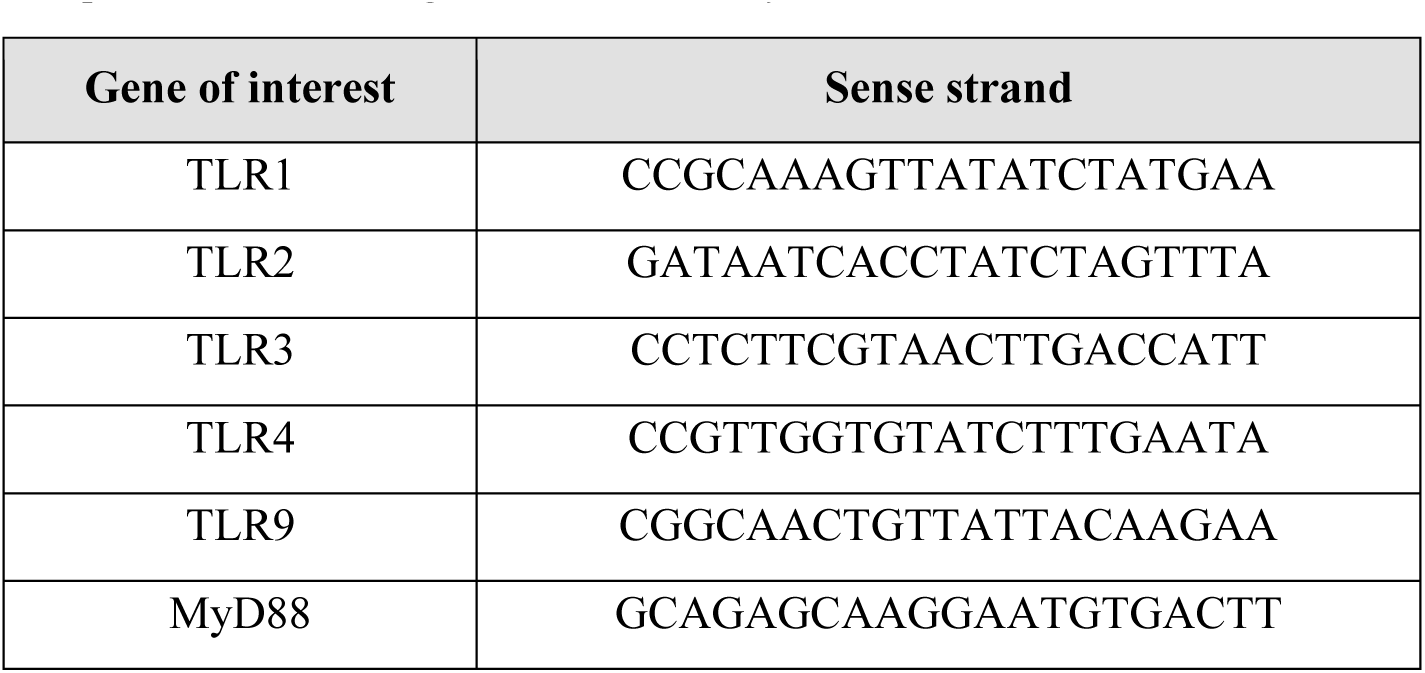
shRNA sequence for TLRs. shRNA sequences were cloned into the pLKO.1 backbone. This table shows the sequences used to target TLR1-9 and MyD88.

All shRNA plasmids with a pLKO.1 background contained Puromycin resistance genes. For co-transfection, the ARF6 shRNA-containing pLKO.1 backbone was modified to replace Puromycin with a Blasticidin resistance gene to enable both plasmids to be individually selected for. To do this, the shARF6-pLKO1 vector was digested using BamHI-HF (NEB #R3136) and KpnI-HF (NEB #R3142) restriction enzymes. The resultant backbone without the Puromycin gene was purified using the QIAquick PCR & Gel Cleanup Kit (Qiagen 28506. A DNA block (Table 3) containing the Blasticidin resistance gene sequence was synthetised (Integrated DNA Technologies) and inserted into the digested backbone using a Gibson cloning reaction.

**Table 3:**
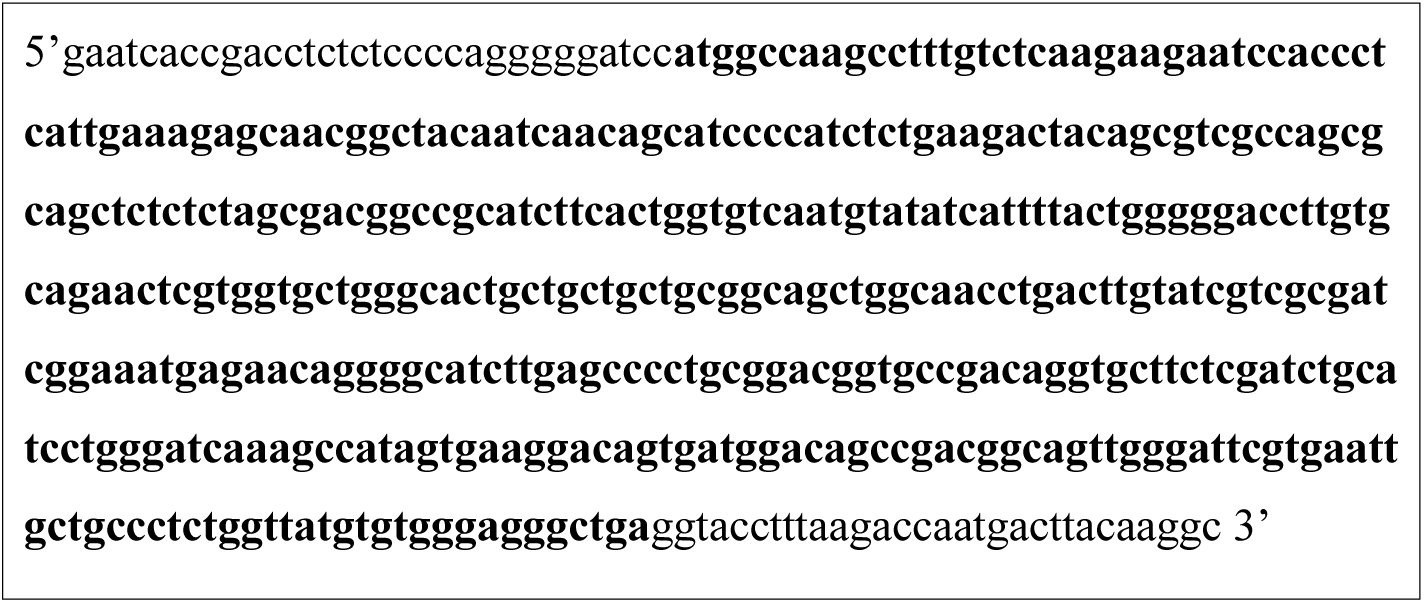
Blasticidin resistance gene sequence. Blasticidin resistance gene was cloned into the pLKO.1 backbone using Gibson cloning. The bold letters represent the Blasticidin resistance gene, which is flanked by Gibson overhangs for integration into pLKO.1 backbone digested using BamHI and KpnI restriction enzymes.

### Lentiviral infection

Lentiviruses for shRNA transduction were generated in HEK293T cells (Takara). 8.4 µg of the vector of interest, 5.6 µg pCMV-dR8.2 dvpr (Addgene #8455) and 4.2 µg pCMV-VSVG (Addgene #8454) were mixed in serum-free OptiMEM media (Gibco) and incubated for 5 min at room temperature. Polyethyleneimine (3 µl/µg of DNA) was mixed in OptiMEM and then the two mixtures combined and incubated at room temperature for 25 min. Following this, the mix was added to the HEK293T cell culture plate dropwise and incubated for 6 hours. Finally, serum-free media was replaced with complete media and cells incubated for 48 hours. Viral supernatant was collected and passed through 0.45 um filters and either directly used in experiments or stored at - 70 °C.

Transduction of MIA PaCa-2 cells using lentiviruses was done in 1:1 ratio of viral supernatant to media and supplemented with 8 µg/ml Polybrene (Sigma-Aldrich #TR-1003-G). Following an overnight incubation, cells were washed with PBS and cultured in complete media containing appropriate selection antibiotics (3 µg/ml of Puromycin or 15 µg/ml of Blasticidin; additionally, prior titration experiments confirmed that with these concentrations, Puromycin selection was completed in 2 days, and Blasticidin in 5-6 days).

### Western Blot

Whole-cell extracts were obtained using an SDS lysis buffer (2% SDS, 50 mM Tris-HCl, and 10% glycerol). Samples were homogenized using the ICOPLUS Insulin Syringe (Twister Medical, N14080). Samples were quantified and mixed with 4× Laemmli sample buffer (Bio-Rad) and boiled at 95 °C for 5 min. Proteins were separated by SDS–polyacrylamide gel electrophoresis and detected with the following antibodies: ARF6 (Santa Cruz Biotechnology sc-7971), ASAP1 (Santa Cruz Biotechnology sc-81896), and Tubulin (Sigma-Aldrich; T6557).

### RNA-seq sample preparation

MIA PaCa-2 cells were treated with shRNAs for ARF6, ASAP1 or control NT (day 0) and followed by Puromycin (3µg/ml, Sigma-Aldrich P8833-25G) selection starting on day 1. Based on titration experiments, Puromycin takes 2 days to kill non-transduced MIA PaCa-2 cells; nevertheless, throughout the entire period of the experiment, cells were maintained in Puromycin to avoid the growth of any remaining non-transduced single cell colonies. On days 3, 7, 14 and 21 cells were trypsinised, centrifuged and cell pellets flash frozen using liquid nitrogen. Storage of cell pellets was at -70 °C.

Once samples from all time points were collected, RNA extraction was performed using the Invitrogen PureLink RNA Mini Kit (Thermo Fisher Scientific #12183018A), as per the manufacturer’s protocol. Sequencing libraries were prepared using the poly(A) selection method and sequenced on an Illumina HiSeq 2500 system at the CRG Genomics Core Facility. Samples were sequenced using 3 biological replicates for each time point and treatment.

### RNA-seq Analysis

Single-end, 50-bp-long reads were aligned to the GRCh37.p13 Homo Sapiens reference genome using the STAR Aligner (version 2.7.6a).^88^ Gene level counts were obtained by STAR using the --quantMode GeneCounts option, using gene annotations downloaded from Gencode (Release 19 GRCh37.p13). Differential expression analysis was performed in R (version 4.0.2) using the DESeq2 package (version 1.30.0).^89^ Genes with adjusted p-value < 0.05 were considered differentially expressed. The lfcShrink() function from DESeq2 was used for visualization purposes. Gene Set Enrichment Analysis (GSEA) was performed using the clusterProfiler package (version 3.18.0)^90^ in R (version 4.0.2). PROGENy analysis was performed in R (version 4.0.3) using DESeq2 normalised counts data as input for the progeny package (version 1.12.0).^34^ Network propagation using PropaNet was performed in Python (version 2.7) using DESeq2 normalised counts data, and a network of TFs and differentially expressed target genes using ChEA3 enrichment analysis.^36^ PropaNet ranks TFs using influence maximisation, followed identification of most important TFs through network propagation.

### ATAC-seq sample preparation

MIA PaCa-2 cells were treated with shRNAs for ARF6, ASAP1 or control NT (day 0) and followed by Puromycin (3 µg/ml, Sigma-Aldrich P8833-25G) selection starting on day 1. Throughout the entire period of the experiment, cells were maintained in Puromycin to avoid the growth of any remaining non-transduced single cell colonies. On days 3, 7, 14 and 21 cells were trypsinised, centrifuged and their cell pellets flash frozen using liquid nitrogen. Storage of cell pellets was at -70 °C.

0.5×10^5^ MIA PaCa-2 cells for each treated condition/replicate infected was collected and treated with transposase Tn5 (Nextera Tn5 Transposase; Illumina Cat #FC-121-1030). DNA was purified using AMPure XP beads to remove large fragments (0.5 × beads; >1kb) and small fragments (1.5 × beads; < 100bp). Samples were then amplified using NEBNext high-Fidelity 2x PCR Master Mix (New England Labs #M0541) with primers containing a barcode to generate the libraries, as previously described.^91^ Each condition was amplified using a combination of the forward primer and one of the reverse primers containing the adaptors listed in the primer table below (Table 4), and sequenced accordingly to allow for segregation of reads. The number of cycles of library amplification was calculated as previously described.^92^ DNA was purified using a MinElute PCR Purification Kit (Qiagen), and samples were sequenced using an Illumina HiSeq 2500 at the CRG Genomics Core Facility. Samples were sequenced using 3 biological replicates for each time point and treatment.^92^

**Table 4.**
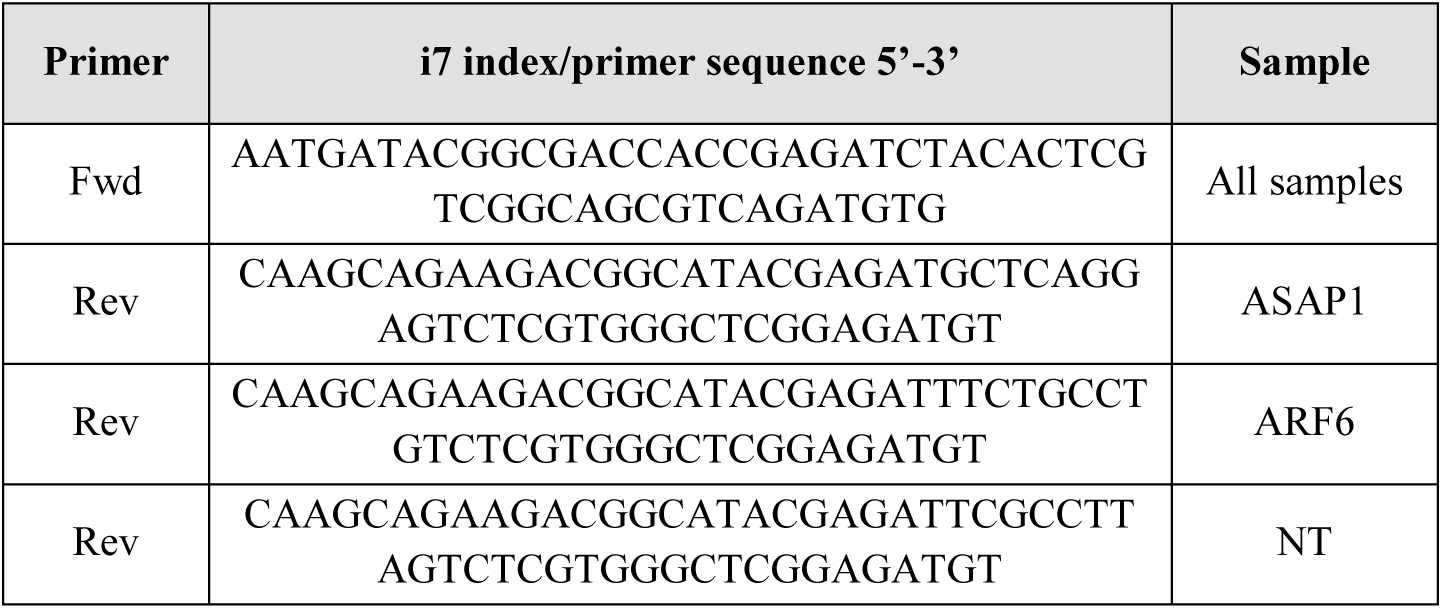
ATAC-seq oligo designs. ATAC-seq primers used for PCR

### ATAC-seq analysis

Paired-end 50 bp reads were adaptor-trimmed using TrimGalore (version 0.6.5). Trimmed reads were aligned to the hg19 genome (UCSC) using BowTie 2 Aligner (version 2.4.2)^93^ with the following parameters: very-sensitive–2000. Aligned reads were filtered using SAM tools (version 1.11)^94^ to retain proper pairs with a MapQ value ≥ 30. Read pairs aligned with ChrM were discarded. Duplicate read pairs were removed using the Picard (version 2.23.8). The read alignment was offset as previously described.^91^ Peaks were called using MACS2 (version 2.2.7.1)^95^ with false discovery rate (FDR) < 0.01. Differentially accessible regions were determined using the DiffBind package (version 3.0.7) in R (version 4.0.2) with false discovery rate (FDR) < 0.005. Genomic annotation of differential regions was performed using HOMER (version 4.11).^96^ Normalised read coverage values were obtained using deepTools (version 3.5.0).^97^ Coverage density heatmaps were generated using deepTools with the option reference point and considering flanking regions 1 kb upstream and downstream from the centre of the peaks.

### Integration of RNA-seq and ATAC-seq

Integration of RNA-seq and ATAC-seq was performed using the normalised log2 gene counts from DESeq2 (see section 3.3.5) and genomic annotation of differential regions from HOMER (see section 3.3.7) respectively. ChEA3 was used to perform TF enrichment from differentially expressed genes, producing a list of enriched TFs (filtered for enrichment score < 100) and their respective list of target genes.^36^ To confirm TF-target gene relationship, TF motif enrichment was performed on the promoter (1kb upstream to 300 bp downstream of the transcription start site) of each target gene using the R package MotifDb (version 1.42.0).^98^ TF-target genes were then classified into activators and repressors, and filtered for activators based on a positive correlation between TF and target gene expression levels. Selected target genes were then clustered using the R package NbClust (version 3.0.1, method = complete). Using Spearman’s correlation, ATAC-seq differential opening and closing was mapped onto the target gene expression trends, and clusters showing an overall positive correlation were selected. Finally, selected genes were clustered into adaptive or constitutively upregulated gene sets and GO enrichment was performed using the enrichPathway function from the R package ReactomePA (version 1.42.0, pAdjustMethod = “BH”).

### Reverse transcription quantitative real time PCR (RT-qPCR)

RNA extraction was done from cells collected and pelleted immediately post-culture or from liquid nitrogen flash frozen cell pellets using the Invitrogen PureLink RNA Mini Kit (Thermo Fisher Scientific #12183018A), as per the manufacturer’s protocol. cDNA conversion was done using the High-Capacity RNA-to-cDNA Kit (Thermo Fisher Scientific #4387406) following which the Applied Biosystems Power SYBR Green PCR Master Mix was used to perform quantitative PCR with 25 ng of cDNA in 10µl reactions as per manufacturer’s instructions. qPCR reactions were performed using primers designed on primer BLAST (Table 5).^99^ Reactions were performed in triplicate (both biological and technical) in a 384-well plate using the ViiA 7 Real-Time PCR System (Thermo Fisher Scientific) or the 7900HT Fast Real-Time PCR System (Thermo Fisher Scientific).

**Table 5.**
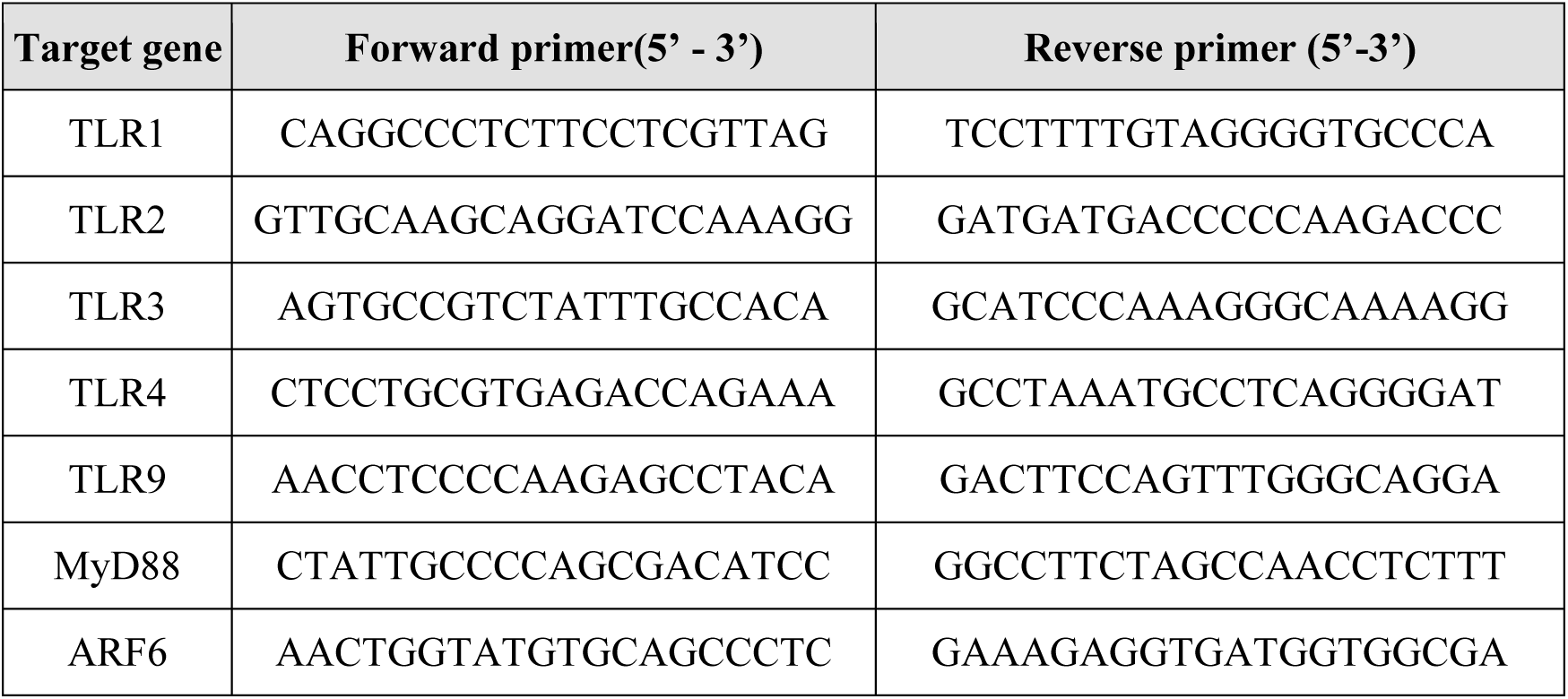
Oligos for qPCR

Quantification of fold change from the qPCR results was performed using the ΔΔCt method.^100^ Cycle threshold (Ct) values were obtained from auto-thresholding that is available in the respective software of the two above mentioned PCR machines. Technical replicate average Ct values of genes of interest (eg: ARF6, TLR1, etc.) were normalised by subtracting the Ct values obtained from the housekeeping gene GAPDH. Next the difference in Ct values was calculated between the control conditions and treatment conditions. Finally, to obtain the relative fold change, the formula 2^-ΔΔCT^ was applied, where the first ΔCT refers to the difference between the gene of interest and housekeeping gene, and the second ΔCT refers to the difference between conditions.

### Transwell invasion assays

MIA PaCa-2 cells were treated with shARF6 or shASAP1 (day 0) and followed by Puromycin (3µg/ml) selection starting on day 1. On days 3, 7, 14 and 21 cells were trypsinised and used for the invasion assays. Assays were performed in 24 well plates containing either 6.5 mm Transwell (8.0 µm pore and polycarbonate membrane, Corning #3422) or 6.5 mm cell culture inserts (8.0 µm pore and polycarbonate membrane, VWR #734-2744). 2 hours prior to the assay, inserts were coated with reconstituted ECM, Matrigel (Growth Factor Reduced (GFR) Basement Membrane Matrix, Corning #354230). To do this, Matrigel stock was diluted to 0.4 µg/ml in ice cold serum-free media and seeded at 100 µl per insert. The insert was then left to incubate at 37 °C for at least 2 hours for gel polymerisation. At the time of seeding, 500 µl of complete DMEM (10% FBS) was added to the lower compartment of each well. Treated MIA PaCa-2 cells were resuspended in serum free media and seeded at 0.6 x 10^5^ per insert.

Cells were allowed to invade for 15 hours following which they were fixed. To do this, inserts were first washed by dipping them in PBS, tapping on a clean paper towel and drying. Using a Q-tip all Matrigel and cells from the inside of the Transwell or insert were removed. Cells were fixed using 4% paraformaldehyde (diluted in PBS) for 15 min at room temperature following which they were washed in PBS and permeabilised using 0.1% Triton-X-100 in 2% BSA/PBS for 30 min. For imaging, nuclei as a proxy for cells, were stained using DAPI (Sigma-Aldrich #MBD0015) for 5 mins at room temperature. Transwell or insert membranes were cut out using a sharp scalpel and mounted on a glass slides using Vectashield Antifade Mounting Medium (Novus Biologicals #H-1000) and sealed using a coverslip and nail polish. Samples were then imaged using the Operetta High Content Screening System (Perkin Elmer) at 20x magnification and at appropriate wavelengths for DAPI (Ex: 405 nm /Em: 461 nm). Since individual images only covered a fraction of the entire membrane, the entire area was captured using multiple images and stitched using the in-built Harmony software (Perkin Elmer).

Post-acquisition image analysis was performed on ImageJ (FIJI).^101^ Briefly, merged images were converted to 8-bit and a threshold applied to it to identify DAPI-stained nuclei. ROIs marking the entire membrane were selected using the elipse selection tool and the mean grey value was measured which quantifies the average cell density. Plotting was done after normalisation to shNT-treated cells.

### Proliferation assay

Proliferation assays were conducted for two experiments. First we used proliferation assays to determine the time-dependent proliferation rate of MIA PaCa-2 cells upon ARF6 or ASAP1 knockdown over a period of 21 days. This was performed using MTT readout. To perform the assay, MIA PaCa-2 cells were treated with shARF6 or shASAP1 (day 0) and followed by Puromycin (3µg/ml) selection starting on day 1. Puromycin takes 2 days to kill untransduced cells. On days 3, 7, 14 and 21 cells were seeded in duplicate for two separate readings. Seeding was done in wells of a 96-well plate in triplicate at 0.1 x 10^5^ cells per well. Cells were allowed to attach and left in the incubators until the following day. Two readings were taken. The first on 1-day-post seeding, and another on 2-day-post seeding. For the readings, cells were incubated for 3 hours at 37 °C with 200 µl 0.5 mg/ml MTT (Panreac AppliChem, #A2231) dissolved in serum-free media. Following the incubation, solubilisation was done in 200 µl of 1:4 DMSO-Isopropanol and the absorbance read at 590nm wavelength. To calculate the proliferation rate, cell readings from 2-days-post seeding were normalised against readings from 1-day-post seeding. This approach removed any technical ambiguities when seeding, and provided directly comparable proliferation rates surrounding each sample collection day.

Second, we used the proliferation assay to evaluate the effect of co-knockdown of ARF6 with TLRs. This was performed using crystal violet staining.^102^ MIA PaCa-2 cells were treated with shRNAs for either ARF6, TLRs or ARF6 and TLRs together. As above, Puromycin (3 µg/ml) or Blasticidin (15 µg/ml) selection was started on day 1 for 6 days. On day 7 the same cell numbers, well-format and time line for measurement was used as in the above MTT-assay. At the time of measurement, cells were first fixed using 4% paraformaldehyde in PBS for 15 min at room temperature. Following PBS washes, cells were incubated in 200 µl Crystal Violet solution for 10 mins, followed by 3-4 PBS washes. Once dry, wells were filled with 200 µl acetic acid for 15 mins at room temperature on a plate shaker to ensure homogenous mixing. Absorbance was measured at 590nm wavelength. Normalisation of readings was done as in the MTT assay to obtain relative growth rates.

### Spheroid formation assay

Spheroid formation assays were performed using the same timeline as in the proliferation assay. MIA PaCa-2 cells were treated with shRNAs for either ARF6, TLRs or ARF6 and TLRs together. As above, Puromycin (3 µg/ml) or Blasticidin (15 µg/ml) selection was started on day 1 for 6 days. On day 7, 800 cells were plated in the Nunclon Sphera Round Bottom 96-well culture plates (Thermofisher Scientific #174925) followed by centrifugation for 4 mins at 800 RPM. For each sample, three technical replicates were seeded. Measurements were taken in the form of images using the Operetta High Content Screening System (Perkin Elmer, see section 3.2.6) at 37 °C and 5% CO_2_. Quantification of spheroid area was done using the in-built Harmony analysis software (Perkin Elmer). As before, cell readings from 2-days-post seeding were normalised against readings from 1-day-post seeding.

### Cell migration assay

Cell migration assays were performed using the same timeline as in the proliferation and spheroid formation assays. MIA PaCa-2 cells were treated with shRNAs for either ARF6, TLRs or ARF6 and TLRs together. As above, Puromycin (3 µg/ml) or Blasticidin (15 µg/ml) selection was started on day 1 for 6 days. On day 7, 0.1 x 10^5^ cells were seeded per well of a 96-well plate. Following 8 hours to allow cells to become adherent, cells were stained with Hoechst 33342 (Life Technologies #H3570) and imaged live for 10 hours using the Operetta High Content Screening System (Perkin Elmer, see section 3.2.6) at 37 °C and 5% CO_2_ at intervals of 20 mins. Post imaging, cell tracking and quantification of distance and displacement per cell was calculated using the in-built Harmony analysis software (Perkin Elmer).

### Mouse tumour growth experiment

In vivo tumour proliferation experiments were performed using subcutaneous injection of MIA PaCa-2 cells in immuno-deficient mice. To do this, we first created a stable MIA PaCa-2 cell line constitutively expressing luciferase. This was done using lentiviral transduction containing the plasmid pLEX-hFL2iG (kindly shared with us by the Antoni Celià-Terrassa group, IMIM), which contains a CMV promoter driven Luciferase-GFP fusion. GFP positive cells were then selected using fluorescence-activated cell sorting (FACS) to identify a luciferase expressing positive population (Figure 3.3.1). FACS was performed on the BDInflux (BD Biosciences) at the CRG/UPF Flow Cytometry Unit.

Prior to injection into mice, MIA PaCa-2^GFP+^ cells were treated with shRNA for either ARF6, TLR2 or ARF6 and TLR2. A non-targeting shNT (Sigma-Aldrich Shc002) control was also used. Following treatment, Puromycin (3 µg/ml) or Blasticidin (15 µg/ml) selection was performed for 6 days, as discussed above. On day 7, cells were resuspended in ice cold PBS at the concentration of 1 x 10^7^ cells/ml.

For injections, 6-week old Athymic Nude mouse, Crl:NU(NCr)-Foxn1nu (Charles River Laboratories) were used. All animal experiments are performed according to the ethical regulations regarding animal research and were conducted under the approval of the Ethics Committee for Animal Experiments (CEEA-PRBB, Barcelona, Spain). Mice were anesthetised using isofluorane and injections were performed in two flanks, left and right, with 1 x 10^6^ cells injected per flank (2 tumours per mice). Tumours were grown for approximately 6 weeks and tumour volume measured from week 2 onwards. We did not observe any visible or quantifiable tumours within the first two weeks, with the exception of the shNT control. At the end of the experiment (6-weeks post injection), luciferase photoluminescence readings were taken for each tumour after administering a peritoneal injection of 100ul D-luciferin potassium salt (Cymit qumica, CAS115144-35-9) in each mice. 8-10 mins were allowed before image acquisition using the IVIS (Perkin Elmer). Analysis of luminescence was done using the Living Image programme (Perkin Elmer). Finally, tumours were harvested, imaged and measured at the endpoint.

### Zebrafish metastasis experiment

In vivo cancer cell metastasis experiments were performed using MIA PaCa-2 injections in zebrafish. To perform these in vivo experiments we first created a constitutively GFP expressing MIA PaCa-2 cell line. To achieve this we transduced MIA PaCa-2 cells with the plasmid lenti-pCMV-KOZAK-EGFP which was kindly shared with us by the Jorge Ferrer group at the CRG, and which contains a CMV promoter-driven enhanced GFP protein inside a lentiviral vector backbone. Cells were selected as single clones using FACS (Supplementary Figure S7 A) performed on the BDInflux (BD Biosciences) at the CRG/UPF Flow Cytometry Unit.

To perform the experiment (Figure 7 A), MIA PaCa-2 GFP+ cells were treated with lentiviruses containing shRNAs for either ARF6, TLR2 or ARF6 and TLR2 together. A non-targeting shNT (Sigma-Aldrich Shc002) control was also used. On the following day, day 1, Puromycin (3 µg/ml) or Blasticidin (15 µg/ml) selection was started for 6 days. On day 7, cells were trypsinised and resuspended in ice cold PBS at 0.1 x 10^6^ cells/µl. These cells were then strained using 40 µm cell strainers (pluriSelect, Germany, SKU 43-10040-40) to obtain a single cell suspension.

For injections, the transgenic zebrafish line Tg(fli:GAL4; UAS:RFP) was used, which allows for the visualization of the vasculature. The line was shared by the group of Berta Alsina at the UPF. All protocols used have been approved by the Institutional Animal Care and Use Ethic Committee (PRBB-IACUEC) and implemented according to national and European regulations. All experiments were carried out in accordance with the principles of the 3Rs (replacement, reduction, and refinement). After collection of spontaneously fertilised eggs, embryos were kept in E3 medium (5 mM NaCl, 0.17 mM KCl, 0.33 mM CaCl_2_, 0.33 mM MgSO_4_) at 28.5 °C before experiments and staged according to morphological criteria^103^ and days-post fertilisation. Starting from the shield stage, 6 hours post-fertilisation, 200 µM PTU (N-phenylthiourea, Sigma-Aldrich) was added to the E3 medium to prevent skin pigmentation. Unfertilised embryos and undeveloped larvae were removed daily.

At 2-dpf, RFP-positive larvae were selected and anesthetised with Tricaine / MS-222 (Sigma-Aldrich). The glass needles necessary for injections were made of borosilicate (GB100T-8P, Science Product) and prepared using a pipette puller (P-97, Sutter Instruments). The needles were connected to a Pneumatic Picopump (WPI). All the injections were performed on zebrafish larvae previously sedated with Tricaine and placed on a Petri dish containing a hardened solution of 1% agarose in E3 water. Cancer cells were injected near the duct of Cuvier (Supplementary Figure S7 B) to facilitate the entry into the circulatory system. After the injection, larvae were kept in E3 at 34 °C. Imaging was performed at day 0 and day 1 post-injection using a zebrafish arraying plate (Hashimoto Electronic Industry, HDK-ZFA101-02a) in the Operetta High Content Screening System. Larvae were euthanized with Tricaine at 4 dpf.

Analysis was performed on ImageJ (FIJI).^101^ To calculate circulating cancer cells, a threshold was applied on the GFP channel to identify MIA PaCa-2 cells, and the total area was calculated per animal at 0 hours post-injection, and 24 hours post-injection. This quantification was performed on the entire animal, after exclusion of the cells remaining within the site of injection. Finally, normalisation was performed as a ratio of cells at 24 hours to 0 hours, to account for variability in injection. Analysis of extravasation was performed by manually counting number of extravasation events per animal. This was done by overlaying the GFP channel showing cells and RFP channel showing vasculature and identifying sites of extravasation.

---

**Supplementary Figure S1.**
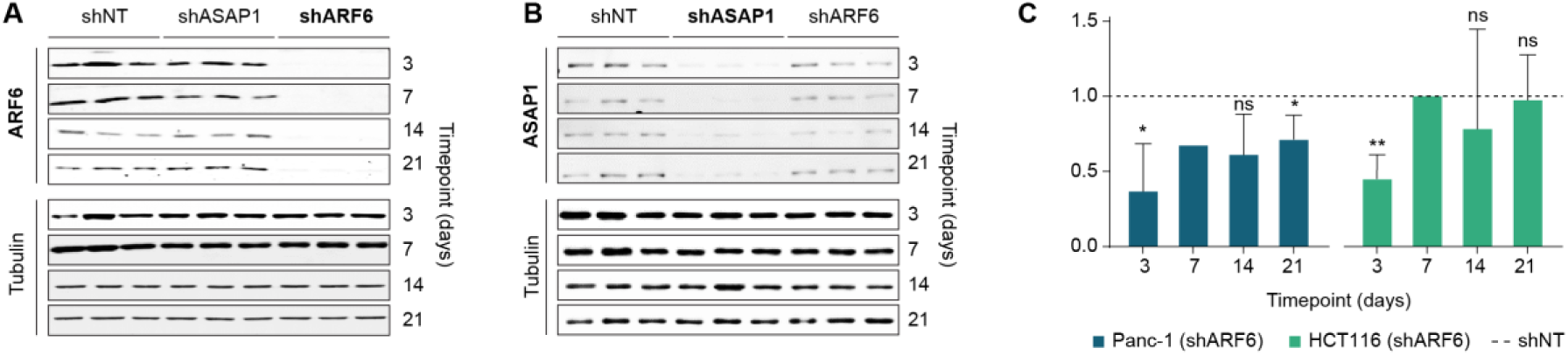
(A) Western Blot validation of samples treated with shRNA for ARF6. The blot shows downregulation of ARF6 only in cells treated with ARF6 shRNA (shARF6), but not in control shNT cells or shASAP1-treated cells. Tubulin was used as a loading control. (B) Western Blot validation of samples treated with shRNA for ASAP1. The blot shows downregulation of ASAP1 only in cells treated with ASAP1 shRNA (shASAP1), but not in control shNT cells or shARF6-treated cells. Tubulin was used as a loading control. (C) Invasion assays in ARF6-depleted Panc-1 and HCT116 cell lines were performed using Transwells coated with Matrigel. Invasion was calculated as number of invaded cells at 15 hours normalised to control. Statistics in *C* are performed using a T-test (* P<0.05, ** P<0.01). Data in *C* represents mean ± sd (n=2).

**Supplementary Figure S2.**
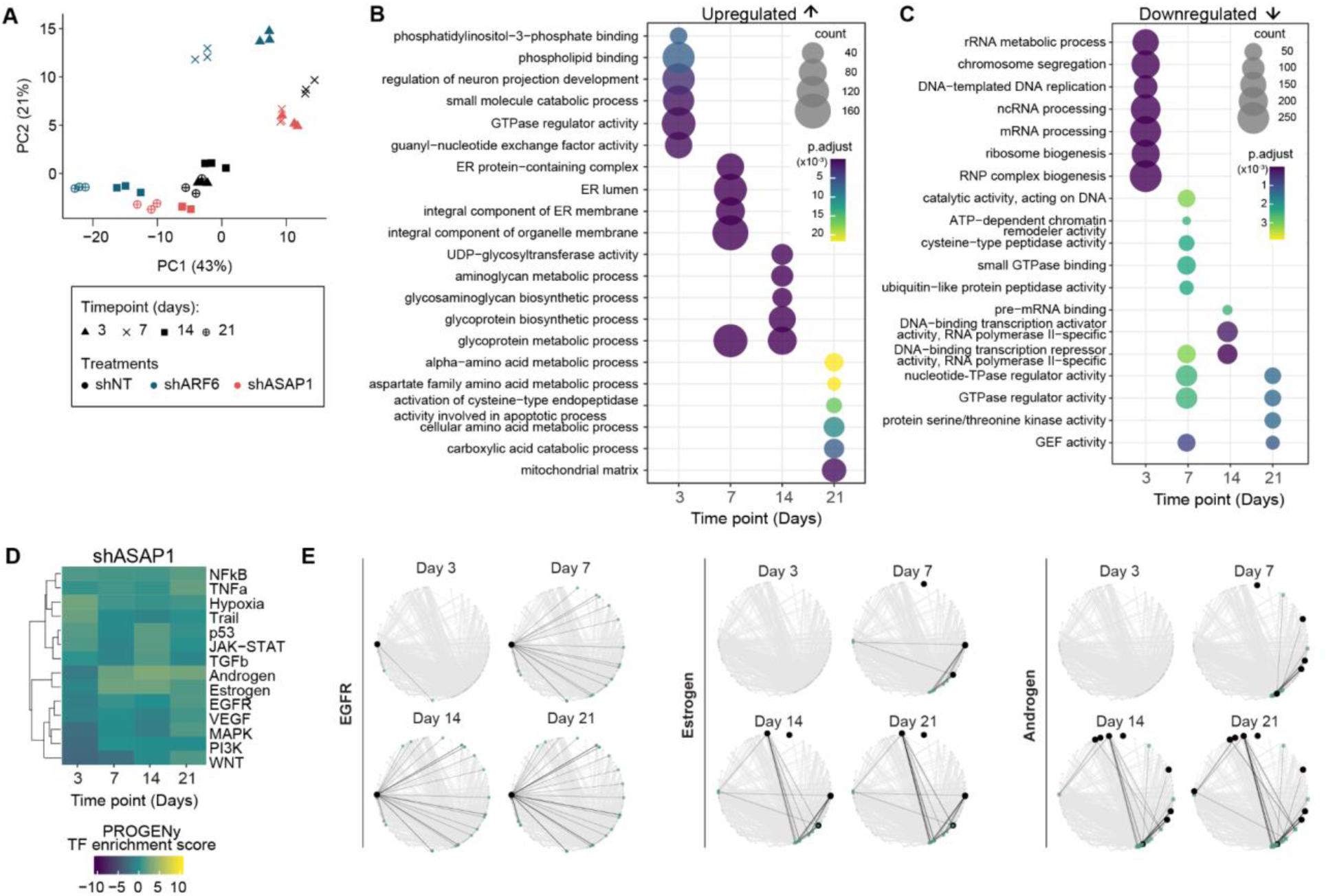
(A) Principal component analysis for transcriptomic data. Sequencing was performed for three biological replicates across four time points, in three different treatments. (B) GO enrichment for upregulated genes in shASAP1 treatment. (C) GO enrichment for downregulated genes in shASAP1 treatment. (D) PROGENy pathway enrichment score from differentially expressed genes in shASAP1-treated cells. (E) PROGENy pathway genes mapped on a PropaNet network created from the enriched TFs and their respective target genes in shARF6-treated cells. Black edges represent all interactions between TFs found in the PROGENy data sets (black nodes) and all their interacting genes (green nodes). See also Figure 2 E.

**Supplementary Figure S3.**
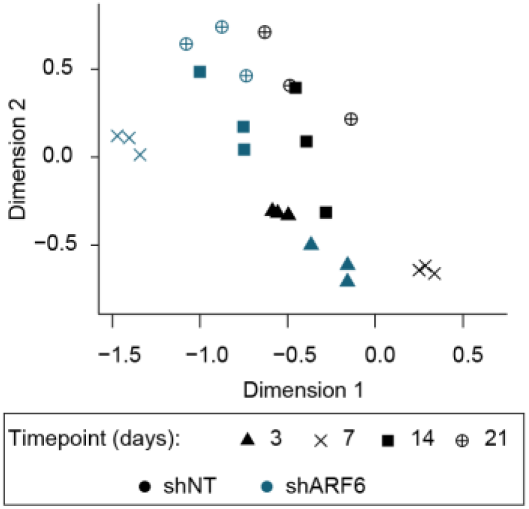
Dimensionality reduction analysis based on ATAC sequencing data. Sequencing was performed for three biological replicates across four time points, in two different treatments.

**Supplementary Figure S4.**
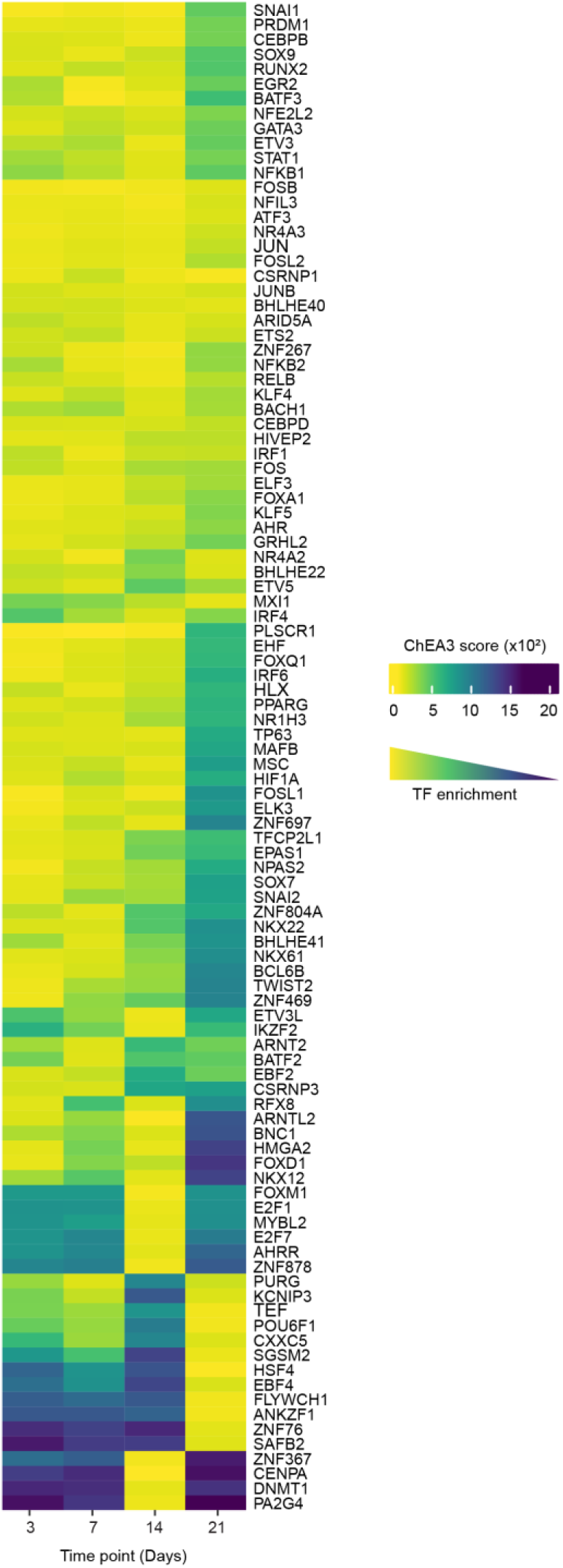
Filtered TFs enriched from ChEA3 analysis of upregulated genes in shARF6-treated MIA PaCa-2 cells. Filtering was done for top TFs (ChEA3 score < 100 for at least one time point).

**Supplementary Figure S5.**
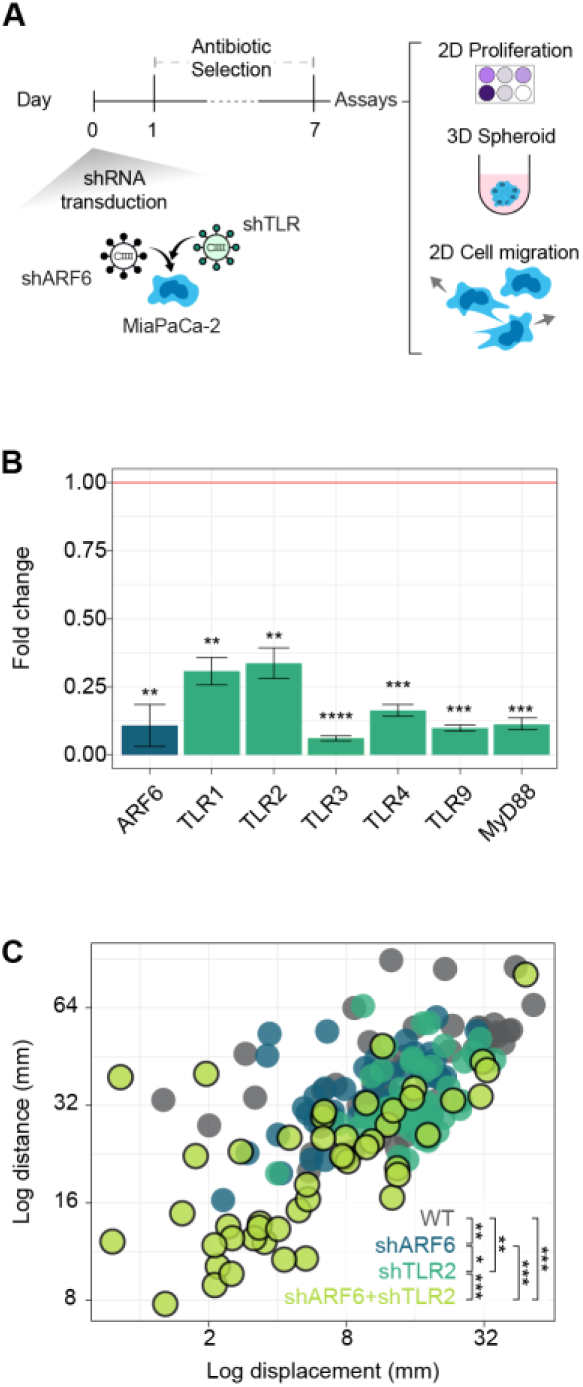
(A) Schematic of experimental strategy to validate the role of TLRs in enabling PDAC adaptation upon ARF6-depletion. MIA PaCa-2 cells were treated with shRNAs for ARF6 and/or TLRs and selected using Puromycin for 7 days following which in vitro assays were performed. (B) Quantitative real-time PCR to validate the downregulation of ARF6, TLRs and MyD88 using shRNAs. Fold change was calculated using the ΔΔCt method (C) Single cell profiles of 50 sampled cells per condition comparing distance and displacement for samples as in Figure 5 D-F. Statistics in *B* were performed using the Wilcoxon test (* P<0.05, ** P<0.01, *** P<0.001, **** P<0.001). Statistics in *C* was performed using the Mann-Whitney-Wilcoxon test (* P<0.05, ** P<0.01, *** P<0.001).

**Supplementary Figure 6.**
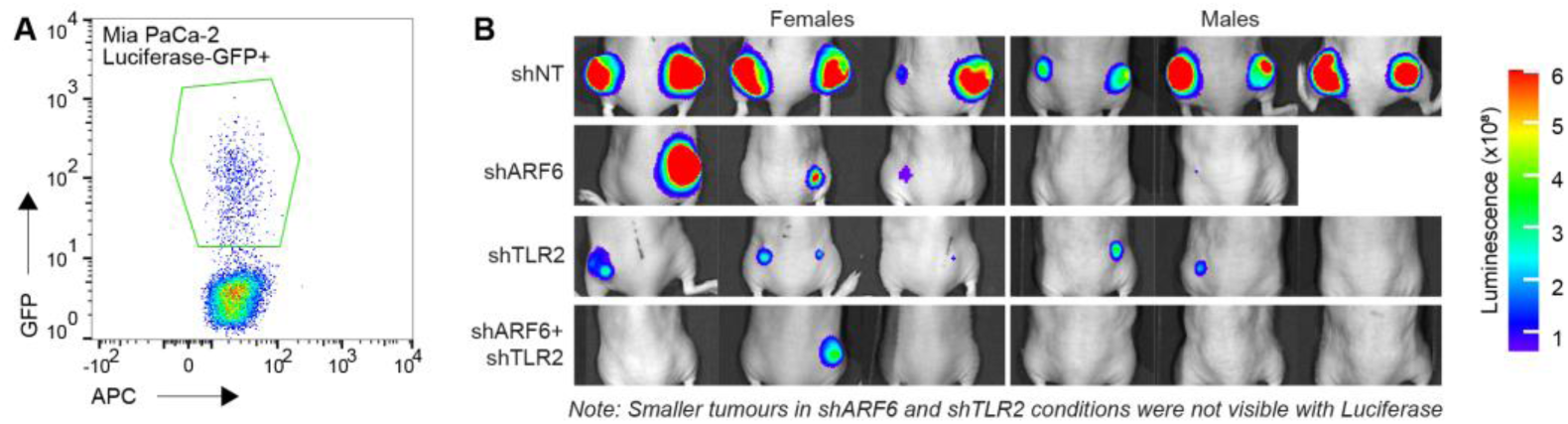
(A) Flow cytometry profile for sorted MIA PaCa-2 cells containing pLEX-hFL2iG, which were selected for GFP+ and used for in vivo mouse xenograft experiments. (B) Luciferase images of tumours just prior to excision (see also Figure 6 B). Small tumours were not visible using Luciferase due to luminescence scale adjustment in comparison to large tumours. This only affected shARF6 and shTLR2 images.

**Supplementary Figure S7.**
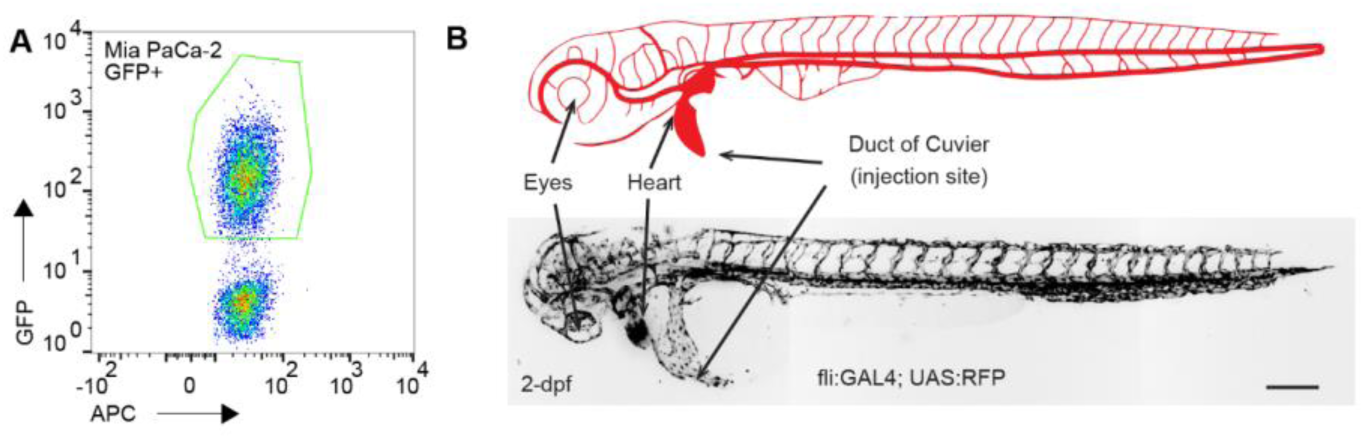
(A) Flow cytometry profile for sorted MIA PaCa-2 cells containing pCMV-KOZAK-EGFP were selected for GFP+ and used for in vivo Zebrafish xenograft experiments. (B) Schematic and confocal microscopy image of zebrafish vasculature. The duct of Cuvier, which is a wide circulation channel connecting the heart and circulatory system, was used as the injection site. Scale bars in *B* represent 200 µm. Schematic in *B* was adapted from Rodel, et. al, 2019.^104^

